# Epigenetic predictors of maximum lifespan and other life history traits in mammals

**DOI:** 10.1101/2021.05.16.444078

**Authors:** C.Z. Li, A. Haghani, T.R. Robeck, D. Villar, A.T. Lu, J. Zhang, C.G. Faulkes, H. Vu, J. Ablaeva, D.M. Adams, R. Ardehali, A. Arneson, C.S. Baker, K. Belov, D.T. Blumstein, E.K. Bors, C.E. Breeze, R.T. Brooke, J.L. Brown, A. Caulton, J.M. Cavin, I. Chatzistamou, H. Chen, P. Chiavellini, O.W. Choi, S. Clarke, J. DeYoung, C.K. Emmons, S. Emmrich, Z. Fei, S.H. Ferguson, C.J. Finno, J.E. Flower, J.M. Gaillard, E. Garde, V.N. Gladyshev, R.G. Goya, M.B. Hanson, M. Haulena, K. Herrick, A.N. Hogan, C.J. Hogg, T.A. Hore, A.J. Jasinska, G. Jones, E. Jourdain, O. Kashpur, H. Katcher, E. Katsumata, V. Kaza, H. Kiaris, M.S. Kobor, P. Kordowitzki, W.R. Koski, B. Larison, S.G. Lee, M. Lehmann, J.F. Lemaitre, A.J. Levine, C. Li, X. Li, D.T.S. Lin, D.M. Lindemann, N. Macoretta, D. Maddox, C.O. Matkin, J.A. Mattison, J. Mergl, J.J. Meudt, G.A. Montano, K. Mozhui, A. Naderi, M. Nagy, P. Narayan, P.W. Nathanielsz, N.B. Nguyen, C. Niehrs, D.T. Odom, A.G. Ophir, E.A. Ostrander, P. O’Tierney Ginn, K.M. Parsons, K.C. Paul, M. Pellegrini, G.M. Pinho, J. Plassais, N.A. Prado, B. Rey, B.R. Ritz, J. Robbins, M. Rodriguez, J. Russell, E. Rydkina, L.L. Sailer, A.B. Salmon, A. Sanghavi, K.M. Schachtschneider, D. Schmitt, L. Schomacher, L.B. Schook, K.E. Sears, A.W. Seifert, A.V. Shindyapina, K. Singh, I. Sinha, R.G. Snell, E. Soltanmohammadi, M.L. Spangler, M. Spriggs, K.J. Steinman, V.J. Sugrue, B. Szladovits, M. Takasugi, E.C. Teeling, B. Van Bonn, S.C. Vernes, H.V. Vinters, M.C. Wallingford, N. Wang, G.S. Wilkinson, R.W. Williams, X.W. Yang, B.G. Young, B. Zhang, Z. Zhang, P. Zhao, Y. Zhao, W. Zhou, J.A. Zoller, J. Ernst, A. Seluanov, K. Raj, V. Gorbunova, S. Horvath, MAMMALIAN METHYLATION CONSORTIUM

## Abstract

Maximum lifespan of a species is the oldest that individuals can survive, reflecting the genetic limit of longevity in an ideal environment. Here we report methylation-based models that accurately predict maximum lifespan (r=0.89), gestational time (r=0.96), and age at sexual maturity (r=0.87), using cytosine methylation patterns collected from over 12,000 samples derived from 192 mammalian species. Our epigenetic maximum lifespan predictor corroborated the extended lifespan in growth hormone receptor knockout mice and rapamycin treated mice. Across dog breeds, epigenetic maximum lifespan correlates positively with breed lifespan but negatively with breed size. Lifespan-related cytosines are located in transcriptional regulatory regions, such as bivalent chromatin promoters and polycomb-repressed regions, which were hypomethylated in long-lived species. The epigenetic estimators of maximum lifespan and other life history traits will be useful for characterizing understudied species and for identifying interventions that extend lifespan.

## Introduction

The maximum lifespan of humans and other mammals appears to be fixed and subject to natural constraints ^1^. The molecular mechanisms underlying these constraints remain poorly understood ^2,3^, despite prior studies correlating maximum lifespan with specific molecular processes and life history strategies ^4–6^. Many authors have suggested that epigenetic mechanisms may play a role in controlling lifespan and aging ^7–15^. The role of epigenetics in mammalian aging is underscored by recent studies demonstrating age reversal through (transient) epigenetic reprogramming with Yamanaka factors ^16–21^.

Here, we have uncovered epigenetic underpinnings of maximum mammalian lifespan and other life history traits using DNA methylation profiles from 192 mammalian species, from 21 taxonomic orders including primates, rodents, bats, cetaceans, and marsupials. Our analyses identified innate cytosine methylation signatures, set at birth, which may predict the maximum lifespan across different species. As such, the methylation levels of these cytosines may not necessarily change across the lifetime of the individual and might be characterized at any point in the lifetime. This contrasts with age-related cytosines, whose levels change predictably with age. We successfully developed methylation-based predictors of time-related life history traits: maximum lifespan, gestation time, and age at sexual maturity across therian mammalian species. In addition to accurately predicting the lifespan of a species, the lifespan predictor correlated with upper limits of lifespan across dog breeds and it corroborated the extended lifespan of growth hormone receptor knockout (dwarf) mice, and mice treated with rapamycin.

## RESULTS

### Large-scale analysis of methylation across mammalian space

To rigorously test whether methylation levels predict maximum lifespan in mammals, we generated cytosine methylation profiles from over 12,000 samples from 60 different tissue types that were derived from 192 species, representing 21 taxonomic orders of mammals. These were profiled using a mammalian methylation array that measures methylation levels of around 36 thousand individual CpGs that along with a 50 base pair flanking DNA sequence are well conserved among mammalian species ^22^. To minimize individual effects within species, we determined the mean methylation value for each CpG of every species, generating a species-summary data set in which each entry is a species average. We also generated a separate data set whose entries correspond to tissue/species strata.

We employed penalized regression analysis to determine the potential relationship between DNA methylation and species lifespan, gestation time, and age at sexual maturity. These life history traits were taken from the current version of the anAge database (Methods) ^2^. The resulting epigenetic predictors exhibited a high level of accuracy according to leave-one-species-out (LOSO) cross validation. The correlation between predicted and actual log maximum lifespan was r=0.89 (**Fig. 1a & 1b**). The log gestation time predictor had an even higher correlation of r=0.96 (**Fig. 1c**) and the log age at sexual maturity predictor exhibited a correlation of r=0.87 (**Fig. 1d**), respectively. We refer to the predicted maximum lifespan, in units of years, as epigenetic maximum lifespan or DNA methylation maximum lifespan, reflecting its provenance.

**Fig. 1.**
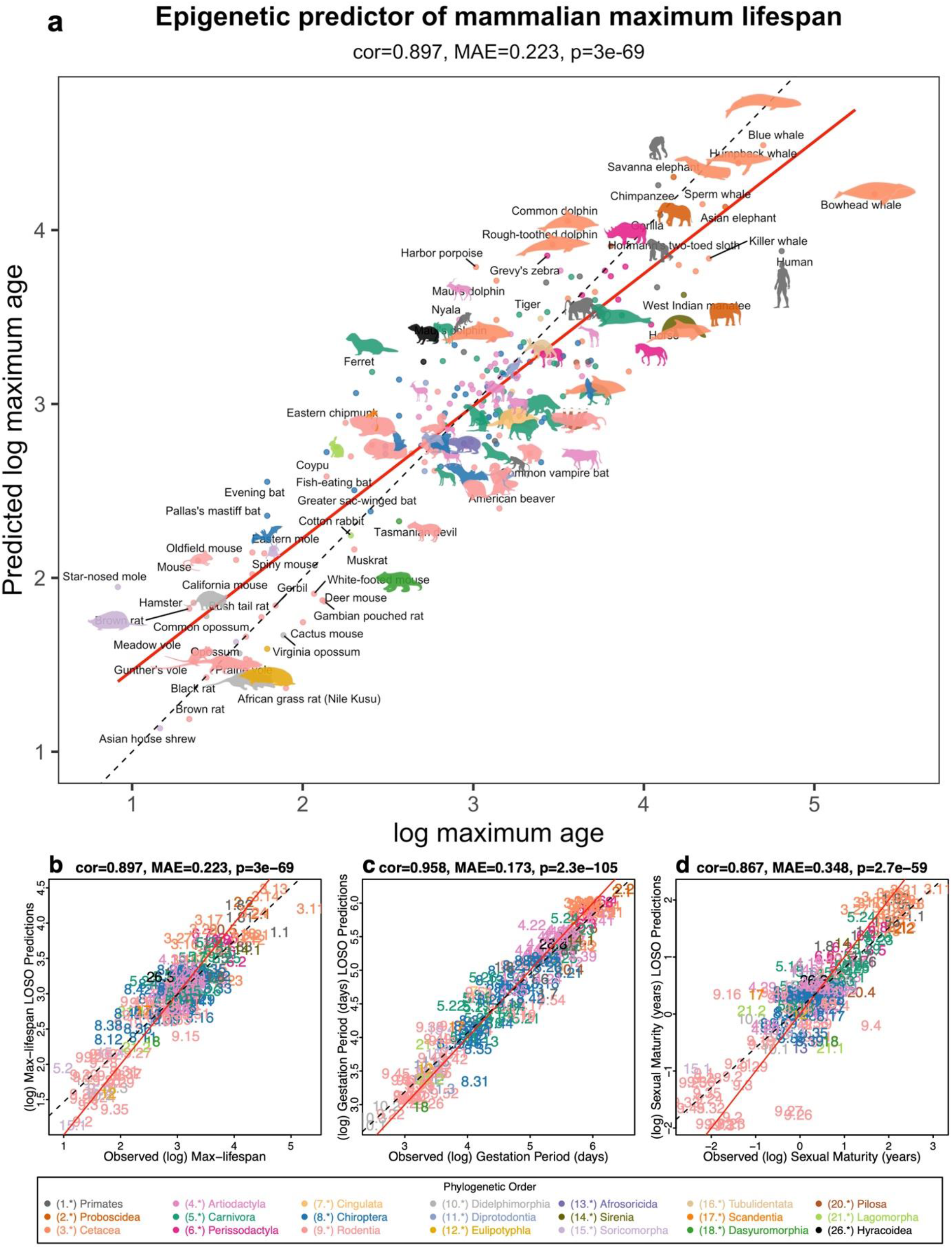
Multivariate predictors of life history traits. Leave one species out (LOSO) cross validation analysis of epigenetic predictors of log (base e) transformed estimates of **a**, maximum lifespan (in years), **b**, gestation time (in days), and **c**, age at sexual maturity (in years). Each species is represented by a number whose integer part denotes the taxonomic order. Each dot corresponds to a different species and is color-coded according to order. Numeric values can be found in **Extended Data Table 1.3**. The titles of the panels report Pearson correlation coefficients, p-values, and median absolute errors (MAE).

To address the concern that epigenetic maximum lifespan might have been confounded with chronological age, we carried out two analyses. First, we developed a second maximum lifespan predictor on the basis of tissue samples from animals younger than their species average age of sexual maturity. **Extended Data Fig. S2** depicts that we are still able to build a good lifespan predictor (r=0.79) based solely on samples from young individuals, even though there were fewer species (n=99) following omission of those from older animals. Second, we show that the predicted maximum lifespan, referred to as epigenetic maximum lifespan, does not correlate with chronological age (**Extended Data Fig. S16**). For most species, the maximum lifespan predicted using female tissues is similar to that based on male tissues (**Extended Data Table 1.3**).

In line with a recent analysis of age- and sex-specific mortality trajectories across mammals ^23,24^, the frequency of species for which females outlive males is about twice as high as for species where the opposite pattern is observed (**Extended Data Table 1.3**). Indeed, females are predicted to have a longer maximum lifespan than males for humans (p=1.E-37, two sample T-test), naked mole rat (p=6.1E-11), vervet monkey (p=0.00012), brown rat (p=0.001), domestic pig (p=0.00014), noctule (p=0.00026), and wapiti elk (p=0.00073). Conversely, males are predicted to live longer than females for Damaraland mole rats (p=0.0027), Seba’s short-tailed bats (p=0.016), Tasmanian devil (p=0.022), and sheep (p=0.00089).

### Cerebellum methylation patterns over-estimate max lifespan

Our predictors of life history traits (including maximum lifespan) were constructed to be agnostic to the underlying tissue. Thus, these predictors should work for DNA derived from any tissue such as blood, skin, liver or brain. As a sensitivity analysis, we compared the predicted values derived from different tissue types. Most tissues showed consistent and comparable predictions, which indicates that these predictors are indeed largely independent of tissue type (**Extended Data Fig. S1** and **Extended Data Fig. S15**). There was however, one notable exception: the methylation profile in the cerebellum leads to substantially higher estimates of maximum lifespan in several different species (**Extended Data Fig. S15 b–g**). For example, the epigenetic maximum lifespan of cerebellar tissue is 491 years for humans, 7 years for mice, 210 years for naked mole-rats, and 85 years for horses, while estimates in non-cerebellar tissue are closer to their actual values: 106 years for humans, 4 years for mice, 34 years for naked mole-rats, and 52 years for horses.

### Lifespan prediction across dog breeds

We conducted two further analyses to evaluate the applicability of the mammalian maximum lifespan predictor to dog breeds with greatly varying lifespans.

We applied it to n=565 blood samples from 51 different breeds ^25^. We averaged the predicted maximum lifespan, i.e. the epigenetic maximum age, by dog breed. We found a positive correlation (r=0.31, p=0.029, **Fig. 2a**) between the average predicted value (epigenetic maximum lifespan) and the upper limit of life expectancy estimated by the American Kennel Club and other registered bodies ^26^. Since the latter may be sub-optimal for ascertaining maximum lifespan of breeds, we employed, instead, adult breed body mass as a correlate of maximum lifespan. We observed a statistically significant negative correlation between epigenetic maximum lifespan (i.e. the predicted value) and average adult breed weight (r=-0.40, p=0.004, **Fig. 2b**). This observation is consistent with the well-attested fact that bigger dogs tend to live shorter lives ^27^.

**Fig. 2.**
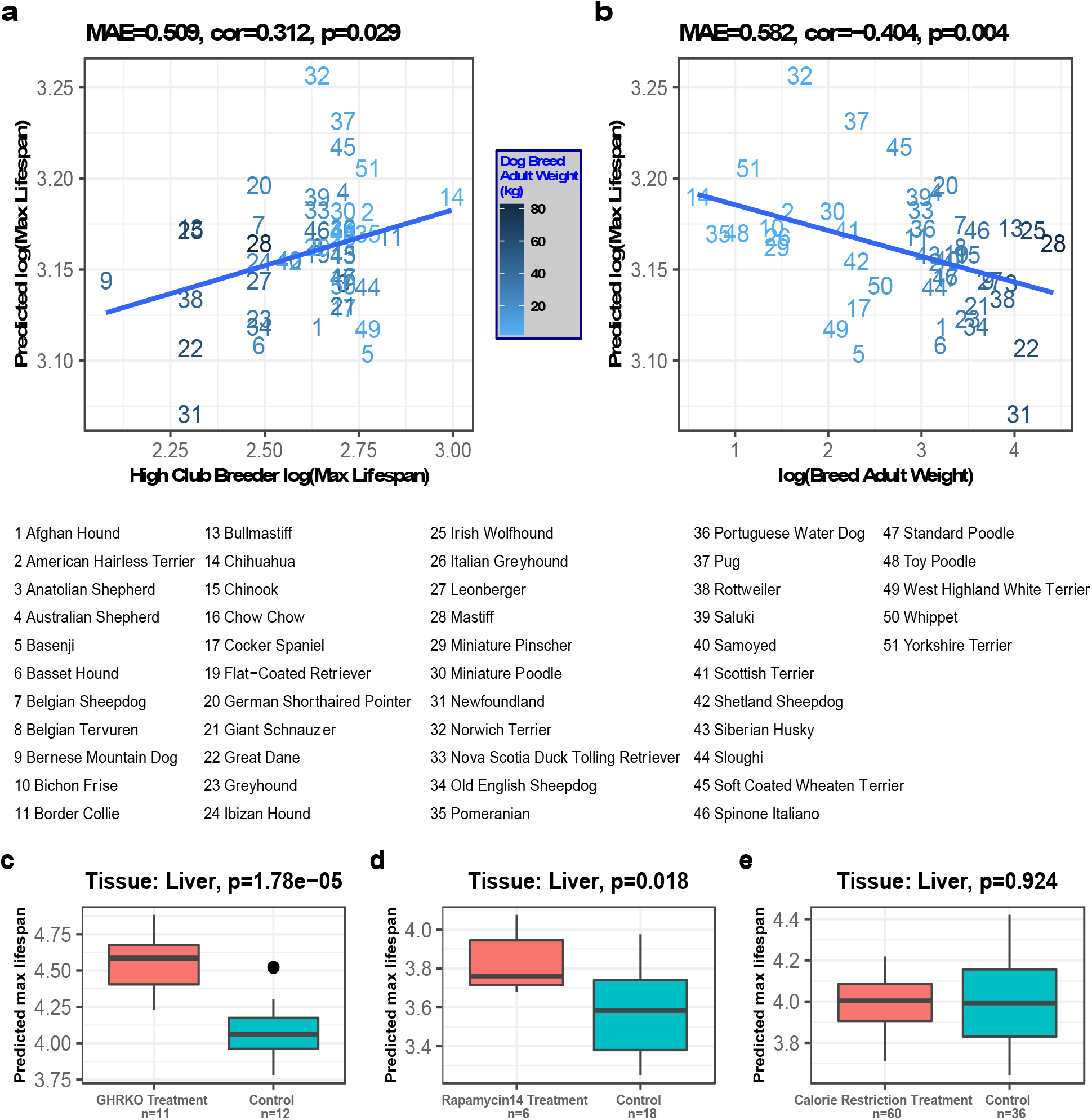
Dog breeds and murine interventions. **a,b**, Log transformed value of epigenetic maximum lifespans (y-axis) in 51 dog breeds based on n=565 blood samples. Each dot (dog breed) represents the average value across multiple blood samples from the same breed. **a**, upper limit of the breed lifespan according to club breeders’ estimate (x-axis), **b**, log transformed value of the average adult weight of the breed. Murine studies of **c**, growth hormone receptor knockout, **d**, rapamycin treatment, and **e**, caloric restriction. Epigenetic maximum lifespan (y-axis) versus group status. The title reports the tissue type (liver) and a Student t-test p-value. Group sizes are reported under the group labels (x-axis).

### Lifespan prediction in murine studies

We further tested the performance of the maximum lifespan predictor in three experimental perturbation set-ups associated with an increase in lifespan. First, we used it to epigenetically predict the lifespan of growth hormone receptor knockout (“dwarf”) mice which are known to have an increased maximum lifespan ^28^. Consistently, the maximum lifespan predictor estimated a longer epigenetic maximum age for dwarf mice than that of regular sized control mice (Student T-test p=1.78e-5)(**Fig. 2c**).

Next, we evaluated the effect of rapamycin in mice since this treatment was reported to extend the life expectancy of model organisms, including rodents (reviewed in ^29^). Using liver samples, we did indeed observe a nominally significant increase in epigenetic maximum lifespan in rapamycin-treated mice (Student T test p=0.018, **Fig. 2d**). However, caloric restriction did not affect the epigenetic maximum lifespan of mice (**Fig. 2e**).

### EWAS of maximum lifespan

To gain insight into biological mechanisms underlying the association between CpG methylation and maximum lifespan it is necessary to identify the precise CpGs whose methylation is associated with lifespan, and to identify genes that are close to them. To this end, we carried out multiple epigenome-wide association studies (EWAS) that differ by how potential confounders were controlled for (average adult body weight, phylogeny or both). We report results from four EWAS studies: (1) Lifespan; a direct regression analysis of lifespan. (2) Weight-adjusted lifespan (AdjWeight); a regression analysis of maximum lifespan after adjustment for average adult species weight. This identifies lifespan-related CpGs that are independent of the body size/weight of the species. (3) Phylogenetic-adjusted lifespan (AdjPhylo); a phylogenetic regression model ^30^ of lifespan, which adjusts for evolutionary relationships between species. (4) Phylogeny and Weight-adjusted lifespan (AdjPhyloWeight); a phylogenetic regression of lifespan after adjustment for average adult species weight. These four analyses were carried out on DNA methylation profiles from five tissues for which there were a sufficiently large number of samples (blood, skin, liver, muscle and brain). At a nominal significance of p<10^−4^, meta-analyses of these tissue DNA methylation profiles identified 4990, 3538, 266, and 338 CpGs that are related to lifespan, lifespan (AdjWeight), lifespan (AdjPhylo), and lifespan (AdjPhyloWeight), respectively (**Fig. 3a**). Since a phylogenetic regression model is more conservative than a “marginal” analysis, phylogenetic regression p-values were less significant. Therefore, we used a more relaxed significance threshold (p<0.005) in our enrichment analyses of phylogenetic EWAS results.

**Fig. 3.**
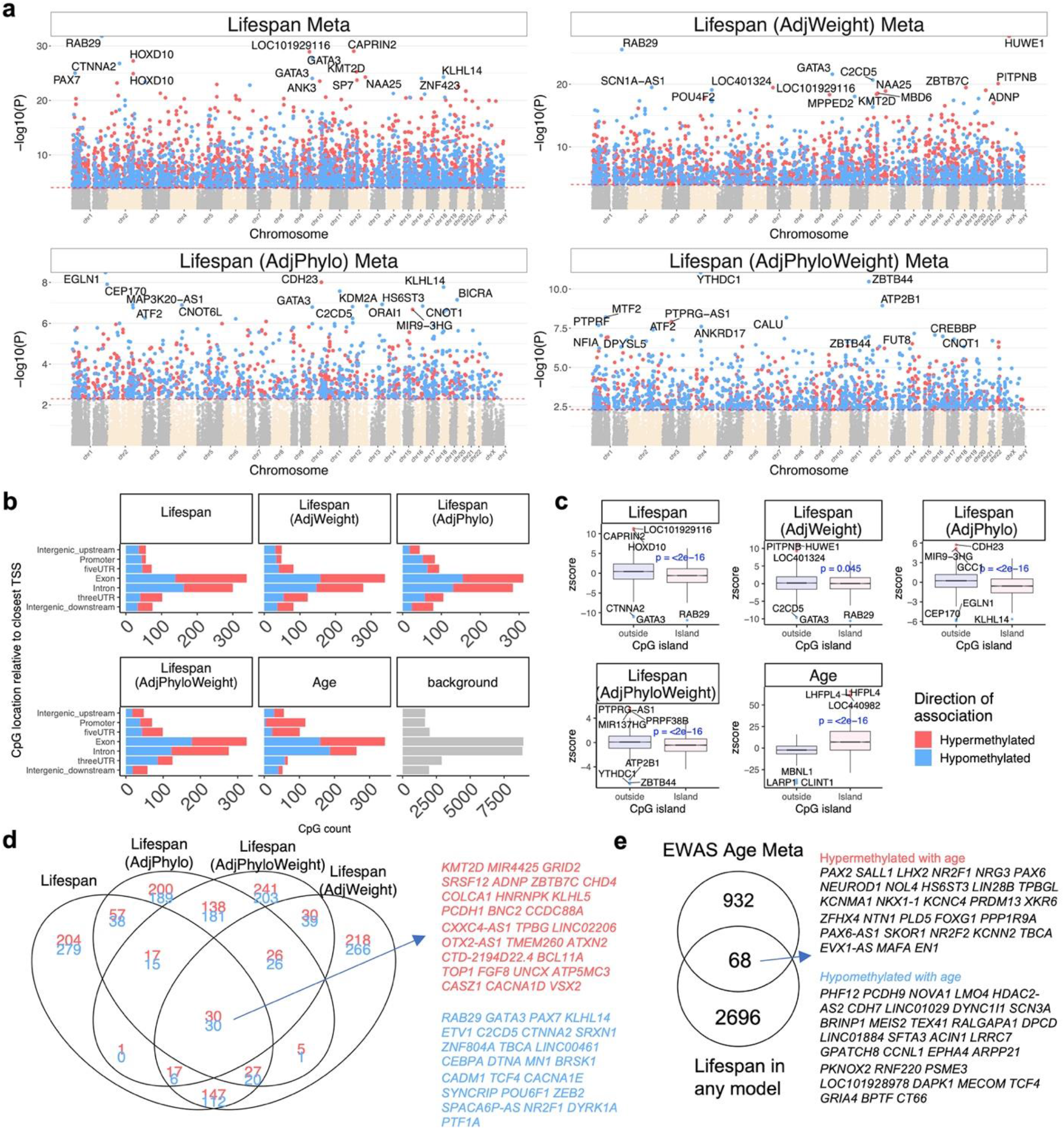
EWAS of mammalian maximum lifespan. The figure represents the meta-analysis (Stouffer’s Z-statistics) of CpG specific association with mammalian maximum lifespan across five tissues: blood, skin, liver, muscle, and brain (all regions). The associations were examined with four different models: 1) lifespan, 2) lifespan adjusted for average adult species weight, 3) lifespan adjusted for phylogenetic relationship from the TimeTree database ^21^, 4) lifespan adjusted for both average weight and phylogeny. **a**, Manhattan plots of EWAS of maximum lifespan in 27966 probes that were experimentally validated to work in both mouse and human genomes. The coordinates are based on the alignment to the Human hg38 genome. The direction of associations with p<10^−4^ (red dotted line) is highlighted by red (positive correlation with maximum lifespan) and blue (negative correlation with maximum lifespan) colors. The phylogenetic regression models were studied at p <0.005. The top 15 CpGs were labeled by the neighboring genes. **b**, Location of the top CpGs in each tissue relative to the closest transcriptional start site. A panel for the top 1000 age related CpGs was added to the figure for comparison ^31^. The grey color in the last panel represents the location of 27966 mammalian BeadChip array probes mapped to the human hg38 genome. **c**, Boxplot of association with mammalian maximum lifespan by human CpG island status. The mean difference was tested by student T test. **d**, Venn diagram of the overlap in the top 1000 (500 per direction) significant CpGs for different models of EWAS of lifespan. The overlap hits were labeled by neighboring genes. **e**, Overlap of CpGs associated with mammalian lifespan and the top 1000 CpGs that relate to chronological age in mammals ^31^. Blood and skin specific results are reported in Extended Data Figures 10-13.

To characterize genes that are potentially involved in determining maximum lifespan, we proceeded to identify those that are proximal to CpGs that are statistically most correlated to lifespan. These are as follows: lifespan was negatively correlated with a CpG in the *RAB29* exon (Z statistic with standard normal distribution under the null hypothesis, z= −11.8 and P=3.9e-32, **Extended Data Fig. S6c**), and positively correlated with a CpG in *CAPRIN2* 3’UTR (z=11.3 and P=1.3e-29, Extended Data Fig. S6a); lifespan (AdjWeight) was positively correlated with a CpG in the *HUWE1* exon (z=11, P=3.8e-28) and negatively correlated with a CpG in *RAB29* exon (z=-10, P=1.5e-23); lifespan (AdjPhylo) was negatively correlated with a CpG in *EGLN1* exon (z=-5.9, P=3.6e-9), and positively correlated with a CpG in *CDH23* intron (z=5.7, P=1.2e-8); and lifespan (AdjPhyloWeight) was negatively correlated with CpGs in *YTHDC1* exon and *ZBTB44* exon. These implicated genes encode proteins that are involved in diverse cellular functions and do not provide an instantly recognizable cellular process or pathway.

We observed that lifespan-related CpGs could be found in genic and intergenic regions with methylation changes in both positive and negative directions. Remarkably, the CpGs in transcription regulatory regions (promoters, 5’UTRs and CpG islands) have positive associations with lifespan (**Fig. 3b**). Interestingly, this pattern contrasts with age-related CpGs, most of which have methylation increasing with age within these regulatory regions (**Fig. 3b**) ^31^. These divergent methylation patterns hint at a possible difference between the process of aging and maximum lifespan. Alternatively, it could be proposed that this difference is complementary whereby the lower methylation baseline of these regulatory regions in long-lived species, provides greater allowance (hence longer time) for accruement of age-related methylation on these promoters. However, CpGs located in 3’UTR, and intergenic regions downstream of gene bodies tended to have positive correlations with lifespan but negative correlations with age (**Fig. 3b**).

Age-related CpGs tend to be different from lifespan-related CpGs. Strikingly, only 68 lifespan-related CpGs from all four models (up to 500 CpGs per direction of association for all of the maximum lifespan meta analyses models) were also among the top 1000 loci that changed with chronological age in mammalian species (**Fig. 3e**). Specifically, the overlap between top 1000 CpG sites from both chronological age EWAS and generic EWAS meta-analysis consists of mere 21 CpG sites, which implies a slight statistical depletion, against the null hypothesis of selecting 1000 CpGs from background at random (hypergeometric test p-value=0.0093, odds ratio=0.60). This suggests that maximum lifespan is largely associated with a stable methylation state of the implicated CpGs, with little change with age throughout the life course of an animal.

As described above, in addition to lifespan, the average adult weight of a species and its phylogenetic position also correlate with DNA methylation. The effects of these were systematically removed in the various EWAS lifespan analyses above. Hence, CpGs that correlate with lifespan in all four analyses can most confidently be assumed to be truly correlated with lifespan. We intersected these CpGs and identified a subset of 60 CpGs with methylation levels that related to lifespan in all four EWAS analyses. Some of these include higher methylation levels in *KMT2D* exon, *MIR4425* intron, *GRID2* intron, and hypomethylation in *RAB29* exon, *GATA3* promoter, and *PAX7* intron (**Fig. 3d**). The identities of these diverse individual genes near the CpGs do not immediately present a hypothesis on how their activities are correlated with lifespan.

Our EWAS meta-analysis combined results from different tissues. As such, we observed a high degree of agreement between the meta-analysis EWAS results and those of tissue-specific EWAS: agreement analyses for generic and phylogenetic EWAS (**Extended Data Fig. S3a-e** and **Extended Data Fig. S4a-e**, respectively). Expectedly, lower agreements are observed between EWAS results from one tissue (e.g. blood) and those from another (e.g. skin, **Extended Data Fig. S3f-o, Extended Data Fig. S4f-o**).

### Gene set enrichment analysis

To uncover biological processes that are potentially linked to lifespan-related CpGs, we employed the Genomic Regions Enrichment of Annotations Tool (GREAT) ^32^ to identify functional annotations associated with genes that are proximal to lifespan related CpGs. Toward this end, we considered the set of top 500 CpGs that have a positive correlation with lifespan (referred to as *lifespan*.*pos* set) and the top 500 CpGs with a negative correlation with lifespan (*lifespan*.*neg set*). We only considered a subset of CpGs by imposing a p-value thresholds, p<1e-4 for generic EWAS and p<5e-3 for phylogenetic EWAS, which resulted in fewer than 500 CpGs in some of the tissue specific analyses.

The resulting analysis provides evidence that these genes were enriched in the following categories: development, metabolism, transcription, immunity, cell proliferation, and cell signaling pathways (**Extended Data Fig. S14**). The predominance of developmental genes and metabolic genes holds true for generic lifespan, WeightAdj-lifespan, PhyloAdj-lifespan and Phylo&WeightAdj-lifespan (**Extended Data Table 3.1.1, 3.1.2, 3.2.1 and 3.2.2** respectively and **Extended Data Fig. S14**).

We related the set of lifespan-related CpGs (up to 500 CpGs) to various human phenotypes previously studied in genome-wide association studies (GWAS). The overlap between lifespan related CpGs and GWAS findings uncovered genes that play a role in neurodegenerative diseases (e.g. hypergeometric test p=0.001 Alzheimer’s disease, p=0.00003 frontotemporal lobar degeneration), body size (e.g. p=0.003 waist to hip ratio), and longevity (e.g. p = 4.37e-6 mother’s attained age, p = 0.0006 longevity > 90, p = 0.003 telomere length, p = 0.007 epigenetic age acceleration of the Hannum clock ^33^) (**Extended Data Fig. S9 and Extended Data Fig. S13**). Some of the differentially methylated genes with variants associated with longevity > 90 years in humans included the *DMRT1* intron, *GPR26* intron, *HOXC5* exon, and *DDX25* exon.

### Chromatin state analysis

To elucidate the genomic context of lifespan-related cytosines, we related them to a universal chromatin state annotation that is based on chromatin marks from over 100 human cell and tissue types ^34^. These chromatin states include those that correspond to constitutive and cell-type-specific activity. Polycomb repressed chromatin states (ReprPC group) are more enriched with positive CpGs (**Fig. 4b**) than with negative CpGs (**Fig. 4a**). Chromatin states of the transcription group (notably Tx7), on the other hand, are enriched with lifespan.neg CpGs after adjusting for weight and phylogeny, especially in blood (p<= 1.4E-14, **Fig. 4a**).

**Fig. 4.**
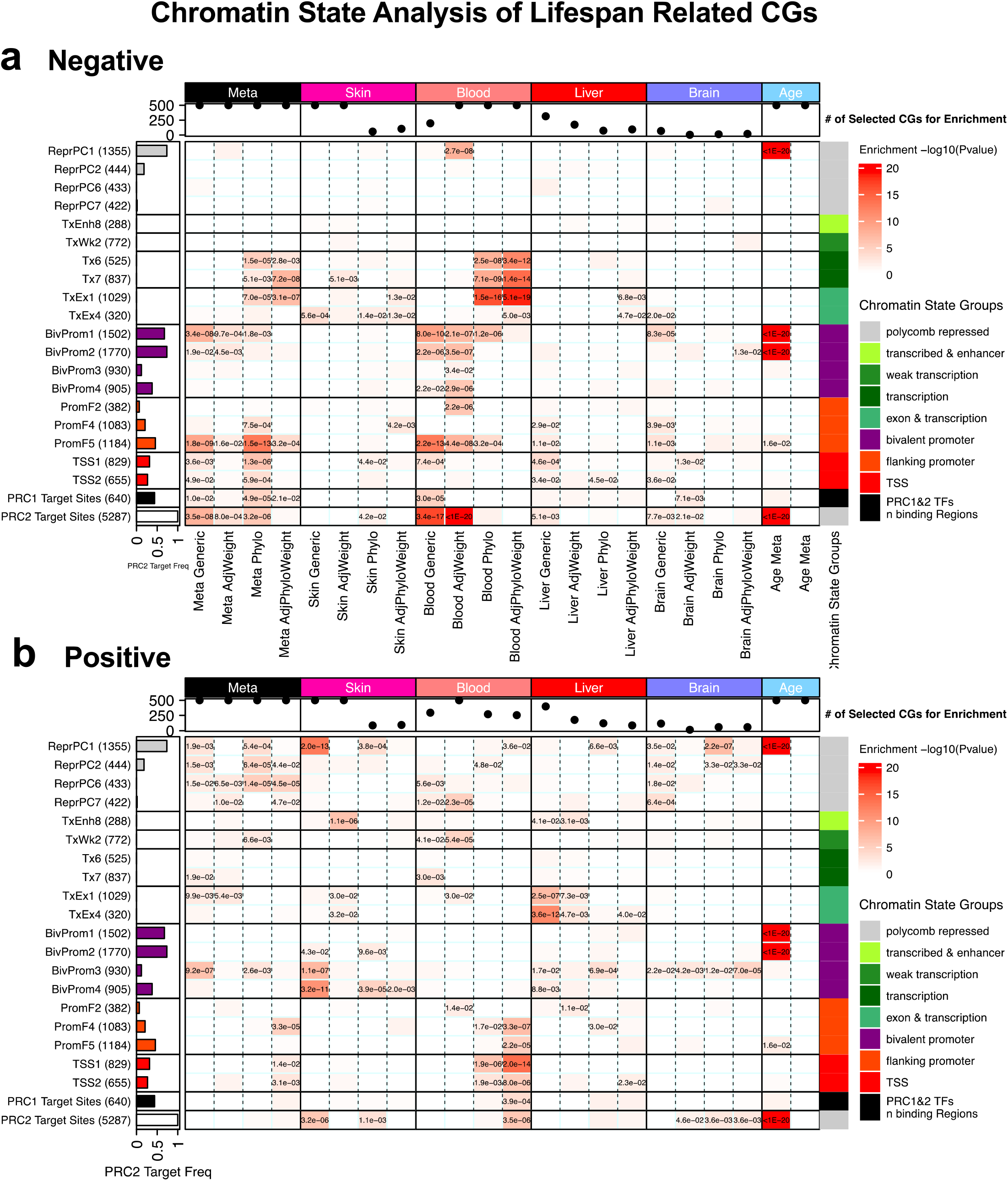
Chromatin State Enrichment analysis of lifespan related CpGs. Enrichment analysis of chromatin states and polycomb repressed complexes with CpGs **a**, that have a negative correlation with maximum lifespan (relatively lower methylation levels in long living species), **b**, that have a positive correlation with maximum lifespan (relatively higher methylation levels long living species). Rows correspond to universal chromatin states ^34^ and are colored and labeled as indicated in the legend on the right. The last two rows correspond to polycomb repressive complex 1 and 2 binding (PRC1 and PRC2, respectively). The columns correspond to sets of CpGs that showed correlations with maximum lifespan in tissue-specific analyses and meta analyses across different tissue types. The figure reports results for generic and phylogenetic regression analysis with and without adjustment for average adult body weight. The last two columns report EWAS results for chronological age (positive and negative association with age, respectively). Row annotations show background CpG frequency for each chromatin state and polycomb repressed complex; column annotations show the number of CpGs selected for each enrichment, up to 500 most significant sites, with a maximum p-value threshold of 0.0001 for generic EWAS and 0.005 for phylogenetic EWAS. Tissues include blood, brain, liver, muscle, and skin. Meta-analysis of considered tissues was conducted via Stouffer’s method, which forms a weighted average of Z-values where the weights are determined by the square root of the sample size.

We observed significant enrichments of lifespan CpGs in bivalent promoter states (BivProm group) but it is important to distinguish positive CpGs, which are enriched in BivProm3 and BivProm4 states, from negative CpGs, which are enriched in BivProm1 and BivProm2 states. Although all are classified as bivalent regions, BivProm3/4 are less bivalent than BivProm1/2 in differentiated cells and exhibit lower occupancy of polycomb repressive complexes PRC1 and PRC2 ^34^. Binding by PRC1 and PRC2 may explain several of the other enrichment results. For example, flanking promoter state PromF5, which is more enriched with lifespan.neg CpGs than other flanking promoter states, also stands out with respect to higher binding of PRC1/2. We also directly conducted enrichment tests for polycomb repressive complexes target sites (PRC1 and PRC2, bottom two rows in **Fig. 4a & 4b**). PRC2 target sites are highly enriched in multiple EWAS analyses, such as weight-adjusted generic EWAS in blood (p<1E-20, **Fig. 4a**).

While PRCs are also strongly implicated in chronological aging, the chromatin states in which maximum lifespan-related CpGs are found do not necessarily overlap with those of chronological age across many species ^31^, as can be seen from the last two columns in **Fig. 4**. CpGs in BivProm1/2 have a negative correlation with maximum lifespan, but gain methylation with age. An exception to this pattern was observed for CpGs in ReprPC1, which exhibit positive correlations with both maximum lifespan and age. Overall, these observations implicate the importance of PRCs in maximum lifespan.

## Discussion

Mammalian species differ dramatically with respect to maximum lifespan. In our data set, maximum mammalian lifespan ranged from 2.5 years (star nosed mole, *Condylura cristata*) to 211 years (bowhead whale, *Balaena mysticetus*). Understanding the mechanisms that are responsible for these vast differences is important for both basic science and clinical translation ^2,35^. Here we demonstrate that DNA methylation of a set of CpG sites can accurately predict species maximum lifespan independent of individual age at sampling time. Our epigenetic predictor of maximum lifespan is largely independent of tissue type with the notable exception of the cerebellum, for which maximum lifespan estimates were substantially over-estimated in several species (e.g. nearly 500 years for humans). This observation, while curious, is consistent with human epigenetic aging clock analyses that estimated the human cerebellum to have the lowest rate of epigenetic aging compared to 30 other tissues ^36^.

The epigenetic predictor of maximum lifespan is expected to have many applications. It can be used to estimate the maximum lifespan of species for which no such metric exists. Although the lifespan predictor is sensitive to some life-extending interventions in mice such as growth hormone receptor knockout or rapamycin treatment, it is not affected by others, including caloric restriction. This, coupled with the fact that predicted maximum lifespan is independent of chronological age (**Extended Data Fig. S16)**, suggest that this predictor is ill-suited for estimating the lifespan of an individual, but rather is useful for estimating the lifespans of species, as it was originally intended. Thus, it will not be suitable for use in most human clinical trials that aim to extend individual human health span. However, we expect that availability of the epigenetic maximum lifespan predictor will usher in a new era of interventional studies that aim to extend the maximum lifespan of a species as a whole.

An important observation that surfaced in this study pertains to the differences between epigenetic maximum lifespan and epigenetic aging. Only 21 CpGs out of the top 1k most significant lifespan related CpGs overlap with the set of top 1k most age-related CpGs. The fact that fewer than expected CpGs overlap (p-value = 0.0093, odds ratio=0.60) reflects in part the statistical approach used to identify lifespan-related CpGs. While there are apparent differences between aging and maximum lifespan at the level of CpGs, there is complementarity at the level of chromatin states. Both lifespan- and age-related CpGs are enriched in bivalent promoters and other chromatin states bound by polycomb repressive complex 2. Interestingly, lifespan-related CpGs in these chromatin states exhibit lower methylation levels in long-lived species than in short lived ones. In other words, it would take longer for these regions to become methylated in long-lived species since the basal methylation levels of these regions are lower. This might be the reason behind their slower physiological rate of aging and longer life. Based on these findings we hypothesize that chromatin organization plays a pivotal role in determining species lifespan. Thus, interventions that enhance chromatin organization and maintenance may increase epigenetic maximum lifespan which is consistent with epigenetic aging theories ^11,19^.

For most species, the epigenetic predictor of lifespan leads to similar results in both sexes. However, we report several species where one sex is predicted to live a longer life than the other sex. Some of these predictions could already be validated based on the existing literature ^23^. While we focused on maximum lifespan, we also present results for correlated life history traits: gestation time and age at sexual maturity.

Overall, these results reveal the importance of epigenetics and cytosine methylation in mammalian lifespan diversity. Indeed, comparative genomics have revealed that the vast majority of genes are conserved between different mammalian species, leading to the postulation that the uniqueness of species is determined by control of regulatory gene expression elements ^37^. This perspective finds support in our analyses where CpGs, which are associated with species lifespan, are enriched in regulatory elements, in terms of genomic position and chromatin state.

## Methods

### Data description

We analyzed methylation data from 192 mammalian species representing 21 different phylogenetic orders (**Extended Data Table 1.1, Fig. 1**). DNA was derived from 64 different tissues and organs including blood, skin, liver, muscle, and several brain regions (**Extended Data Table 1.2**). Materials and Additional File 1 contains details on all the data sets that we have used to conduct analyses. To enhance the reproducibility of our findings we include our updated version of the anAge database ^2^.

### DNA methylation

All data were generated using the mammalian methylation array (HorvathMammalMethylChip40) ^22^. The mammalian methylation array provides high coverage of highly conserved CpGs in mammals. Out of 37,492 CpGs on the array, 35,989 probes were chosen based on high levels of sequence conservation within mammalian species^6^. The particular subset of species each probe is expected to work in is provided in the chip manifest file which can be found at Gene Expression Omnibus (GEO) at NCBI as platform GPL28271 and on our Github webpage. The SeSaMe normalization method was used to define beta values for each probe ^38^. Genome coordinates for different dog breeds have been posted on Github as detailed in the section on Data Availability.

### Multivariate estimators of maximum lifespan

To build a multivariate predictor of maximum lifespan, gestation time, and age at sexual maturity we used elastic net regression ^39^. Since we aimed to build the model on the basis of CpGs that are present/detectable in most species, we restricted the analysis to CpGs with significant median detection p-values (false discovery rate<0.05) ^40^ in 85% of the species. This results in a lower-dimensional dataset consisting of 19779 CpGs. Toward this end, we calculated the median detection p-value (using the SeSaMe normalization method) per species.

We employed two strategies for building lifespan predictors. The first strategy ignored tissue type. Here, all tissue samples from a given species were averaged resulting in a single observation (average) per species. The second strategy formed average values for each stratum defined by tissue type and species. For example, this analysis formed an average value for human blood, human liver, mouse blood, mouse liver, etc. The second approach allowed us to study the influence of tissue type on lifespan predictions. To arrive at unbiased estimates of the predictive accuracy of these lifespan predictors (and other predictors), we used a leave-one-species-out (LOSO) cross validation analysis that iteratively trained the predictive model on all but one species. Next, the predictor was applied to the observations from the left out species. By cycling through the species, we arrived at LOSO estimates for each species. As sensitivity analysis, we also conducted a leave-one-taxonomic-order-out analysis. For example, at one iteration, we trained a model on all available taxonomic orders except for primates. This resulted in findings that were very similar to those from LOSO, e.g. we find a high correlation (r=0.77) between the leave-one-taxonomic-family-out estimate of log maximum lifespan and its actual value.

Considering the high correlation of maximum lifespan and adult weight, we examined the potential confounding effects of species weight on the performance of our model. The LOSO estimate of maximum lifespan was highly correlated (r = 0.60, p=2e-16) with the weight adjusted estimate of log of maximum lifespan. Similarly, a multivariate regression model (dependent variable log of maximum lifespan) revealed that log body weight (Wald test p=0.00045) is a less significant covariate than the estimate of log maximum lifespan (p = 2e-16).

### EWAS of log maximum lifespan

Since the distribution of maximum lifespan was highly skewed, we imposed a log-transformation on maximum lifespan before conducting EWAS. Our EWAS of maximum lifespan focused on 27,966 CpG probes that were experimentally validated to work in both mice and humans^6^. We carried out four types of analyses that differ by how they deal with two potential confounders: adult weight and phylogeny. Our “generic” EWAS corresponds to a marginal correlation analysis where the average methylation level of a given CpG per species was regressed on the (log transformed) maximum lifespan using ordinary least squares regression. The second EWAS approach replaced ordinary least squares regression by phylogenetic regression, the variance-covariance matrix of which modeled evolutionary distances using branch lengths from the Tree of Life web project ^30,41^. To adjust for adult weight, we first regressed log maximum lifespan on log weight and formed residuals. Next the residuals become the dependent variables in the regression models. We carried out EWAS analyses in the following tissues/organs: skin, blood, liver, and brain, based on which we also calculated meta-analysis Z statistics using Stouffer’s method (weighted by square root of corresponding sample sizes). For each of the following 4 approaches of carrying out EWAS enrichment of maximum lifespan, we defined two sets of lifespan related CpGs: lifespan.pos and lifespan.neg consist of up to 500 CpGs each with positive and negative correlations with log maximum lifespan, respectively. We omitted CpGs from these sets of 500 CpGs if their respective p-values exceeded 0.0001 in case of generic EWAS and 0.005 in case of phylogenetic EWAS.

### Genomic region based enrichment studies

We selected up to the top 500 significant CpG, per direction, sites with p-values <1e-4 for generic EWAS and <5e-3 for phylogenetic EWAS. Our genomic region based enrichment analysis used the R package GREAT ^32^ in hg19 assembly. The extension of gene regulatory regions was set at 50 kb and the other options were based on default settings. Since our EWAS focused on 27,966 CpGs that applied to both humans and mice, these probes were used as the background ^22^. By specifying the background, GREAT analysis performed genomic-region based hypergeometric analysis, not confounded by gene sizes and uneven gene coverage.

### EWAS-GWAS based overlap analysis

Our EWAS-GWAS based overlap analysis related the gene sets found by our EWAS of maximum lifespans with the gene sets found by published large-scale GWAS of various phenotypes, across body fat distribution, lipid panel outcomes, metabolic outcomes, neurological diseases, six DNAm based biomarkers, and other age-related traits. Enrichment p values for the overlap between the genes implicated in EWAS and GWAS were based on genomic region based hypergeometric tests as detailed in **Extended Data Tables 3.1.1-3.2.5 and Extended Data Tables 4.1-5.4**.

### Chromatin State Analysis

We conducted chromatin state enrichments using a universal annotation of the human genome annotation that is not specific to one cell or tissue type based on a stacked ChromHMM model recently generated based on over 1000 data sets from diverse human cell and tissue types ^34^. For each EWAS enrichment mentioned above, we utilized a hyper-geometric test to assess significant overlap between chromatin states and the two sets of CpGs that are highly significant in either positive or negative correlations with maximum lifespan. The background set for these hyper-geometric enrichment tests were the 27,966 CpGs that mapped to both human and mouse.

### EWAS of Age

Our meta analysis EWAS of chronological age across different species and tissue types is described in Lu et. al ^31^. EWAS of age was performed on species-tissue strata (approximately 135 strata from 60 species). The meta analysis was carried out in two steps. First, we combined the EWAS results across tissues within the same species to form meta Z scores at species level. Second, we combined the EWAS results across species to form the final meta Z scores for EWAS of age. The background set of CpGs for the enrichment analysis was defined as the 27,966 that map to both mice and humans.

## Conflict of Interest Statement

SH is a founder of the non-profit Epigenetic Clock Development Foundation which plans to license several patents from his employer UC Regents. These patents list SH, AA, and JE as inventors. Robert T. Brooke is the Executive Director of the Epigenetic Clock Development Foundation. The other authors declare no conflicts of interest.

## Data Availability Statement

The data will be made publicly available as part of the data release from the Mammalian Methylation Consortium. Genome annotations of these CpGs can be found on Github https://github.com/shorvath/MammalianMethylationConsortium

## ACKNOWLEDGEMENTS

This work was mainly supported by the Paul G. Allen Frontiers Group (SH). Diego Villar was supported by a personal Fellowship from the British Heart Foundation (FS/18/39/33684). Duncan T Odom was supported by Wellcome (WT202878/Z/16/Z), European Research Council (788937), and Cancer Research UK (20412). Sonja C Verne was supported by a Max Planck Research Group Award from the Max Planck Gesellschaft and a UKRI Future Leaders Fellowship (MR/T021985/1).

