## Extended data figures for "Epigenetic predictors of maximum lifespan and other life history traits in mammals"

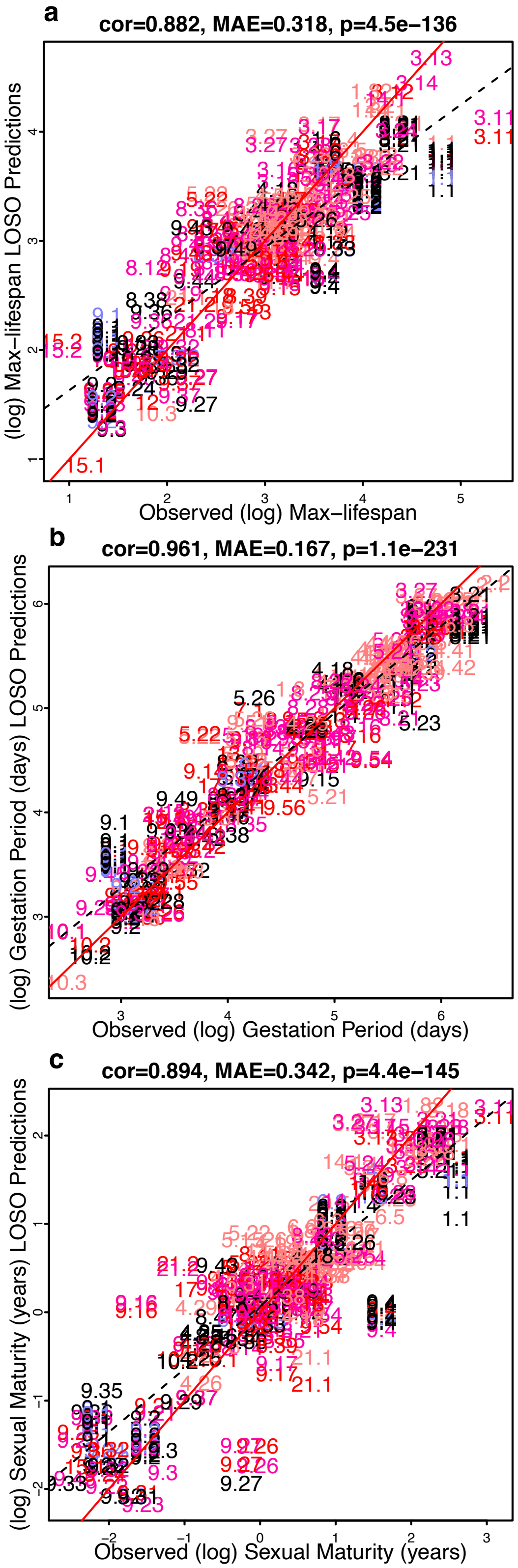


**1 Primates**
1.1 Human
1.2 White−fronted marmoset
1.3 Common marmoset
1.4 Vervet
1.5 Rhesus macaque
1.6 Olive baboon
1.7 Francois leaf monkey
1.8 Red ruffed lemur
1.81 Gorilla
1.82 Chimpanzee
1.83 Orangutan
**2 Proboscidea**
2.1 Asian elephant
2.2 Savanna elephant
**3 Cetacea**
3.11 Bowhead whale
3.12 Sei whale
3.13 Blue whale
3.14 Humpback whale
3.15 Commerson's dolphin
3.16 Maui's dolphin
3.16 Maui's dolphin
3.17 Common dolphin
3.18 Short−finned pilot whale
3.19 Pacific white−sided dolphin
3.21 Killer whale
3.22 Bottlenose dolphin
3.23 Beluga whale
3.24 Sperm whale
3.25 White beaked dolphin
3.26 Sowerbys beaked whale
3.27 Harbor porpoise
3.28 Rough−toothed dolphin
**4 Artiodactyla**
4.11 Springbok
4.12 Cattle
4.13 White−bearded gnu
4.14 Thomson's gazelle
4.15 Sable antelope
4.16 Grant's gazelle
4.17 Soemmerring's gazelle
4.18 Sheep
4.19 Nyala
4.21 Greater kudu
4.22 Roe deer
4.23 Indian muntjac
4.24 White−tailed deer
4.25 Domestic pig
4.26 Miniature pig
4.27 Domestic goat
4.28 Red deer
4.29 Slender Horn gazelle
4.31 Wapity elk
4.32 Roan antelope
4.33 Nile lechwe
4.34 Gerenuk
4.35 Dama gazelle
4.36 Eland
4.37 Bongo
4.38 Lesser kudu
4.39 Alpaca
4.41 Giraffe.reticulata.hybrid
4.42 Okapi
4.43 Addax
4.44 Impala
4.45 Sitatunga
**5 Carnivora**
5.11 Coyote
5.12 Dog
5.13 African wild dog
5.14 Cheetah
5.15 Cat
5.16 Lion
5.17 Tiger
5.18 Meerkat
5.19 Asian small−clawed otter
5.21 Southern sea otter
5.22 Ferret
5.23 Pacific walrus
5.24 California sea lion
5.24 California sea lion
5.25 Harbor seal
5.26 Black bear
5.27 Maned wolf
5.28 Steller sea lion
5.29 Australian sea lion
5.31 Red fox
5.32 Bobcat
5.33 Florida panther
5.34 Spotted hyena
**6 Perissodactyla**
6.2 Horse
6.3 Zebra
6.4 Southern white rhinoceros
6.5 Eastern black rhinoceros
6.6 Somali wild ass
6.7 Grevy's zebra
6.8 Greater one horned rhino
**7 Cingulata**
7.1 Big hairy armadillo
**8 Chiroptera**
8.11 Greater sac−winged bat
8.12 Pallas's mastiff bat
8.13 Mexican free−tailed bat
8.14 Jamaican fruit bat
8.15 Seba's short−tailed bat
8.16 Common vampire bat
8.17 Pale spear−nosed bat
8.18 Greater spear−nosed bat
8.19 Lesser short−nosed fruit bat
8.21 Straw−colored fruit bat
8.22 Indian fruit bat
8.23 Variable flying fox
8.24 Grey−headed flying fox
8.25 Little golden−mantled flying fox
8.26 Rodriguez flying fox
8.27 Large flying fox
8.28 Egyptian fruit bat
8.29 Greater horseshoe bat
8.31 Pallid bat
8.32 Big brown bat
8.33 Brandt's bat
8.34 Little brown bat
8.35 Greater mouse−eared bat
8.36 Fish−eating bat
8.37 Noctule
8.38 Evening bat
8.39 Pipistrelle
8.41 Brown long eared bat
8.42 Lesser long−nosed bat
8.43 Proboscis bat
8.44 Halcyon horseshoe bat
8.45 Noack's roundleaf bat
8.46 Western long−eared myotis
8.47 Fringed myotis
**9 Rodentia**
9.1 Mouse
9.14 Cape mole rat
9.15 American beaver
9.16 Guinea pig
9.17 Capybara
9.18 Chichillah
9.19 Coypu
9.2 Brown rat
9.2 Brown rat
9.21 Mole vole Ellobius
9.22 Hamster
9.23 Gunther's vole
9.24 Prairie vole
9.25 Meadow vole
9.26 Bush tail rat
9.27 Muskrat
9.28 California mouse
9.29 Cactus mouse
9.3 Black rat
9.31 White−footed mouse
9.32 Deer mouse
9.33 Oldfield mouse
9.34 Lowland paca
9.35 African grass rat (Nile Kusu)
9.36 Gerbil
9.36 Gerbil
9.37 Gambian pouched rat
9.38 Yellow−bellied marmot
9.39 Groundhog
9.4 Naked mole rat
9.41 Eastern grey squirrel
9.42 Fox squirrel
9.43 Eastern chipmunk
9.44 Red squirrel
9.45 Chinese hamster
9.49 Blind mole rat
9.51 Crested porcupine
9.52 Spiny mouse
9.54 North American porcupine
9.55 Middle East blind mole rat
9.56 Damaraland mole rat
**10 Didelphimorphia**
10.1 Common opossum
10.2 Opossum
10.3 Virginia opossum
**11 Diprotodontia**
11.1 Tammar wallaby
11.2 Western grey kangaroo
11.3 Eastern grey kangaroo
11.4 Red−necked wallaby
11.5 Red kangaroo
11.6 Agile wallaby
11.7 Hill wallaroo
**12 Eulipotyphla**
12 Spanish mole
**13 Afrosoricida**
13 Lesser hedgehog tenrec
**14 Sirenia**
14.1 West Indian manatee
**15 Soricomorpha**
15.1 Asian house shrew
15.2 Star−nosed mole
15.3 Hairy tail mole
15.4 Eastern mole
**16 Tubulidentata**
16 Aardvark
**17 Scandentia**
17 Northern treeshrew
**18 Dasyuromorphia**
18 Tasmanian devil
**20 Pilosa**
20.4 Linne's Two−toed sloth
20.5 Hoffmann's two−toed sloth
**21 Lagomorpha**
21.1 European rabbit
21.2 Cotton rabbit
**26 Hyracoidea**
26.3 Rock hyrax

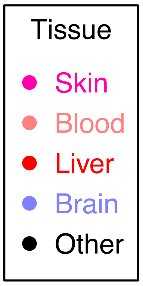

**Extended Data Fig. S1 | Predictors of Species-Tissue Combinations.** A penalized joint linear model used to predict species lifespan (Elastic Net). Same framework as that of **Fig. 1**, except that it distinguishes tissue types. CpG probes are averaged by each species-tissue combination. Different tissues within the same species share the same maximum lifespan, but retain different methylation levels. Three panels show predictors for **a,** maximum lifespan (in years), **b,** gestation time (in days), and **c,** age at sexual maturity (in years). Note: *LOSO, definition deferred to that of **Fig. 1**. Designated Mammalian numbers in scatter plot panels and the figure legend are the same as those of main **Fig. 1.**

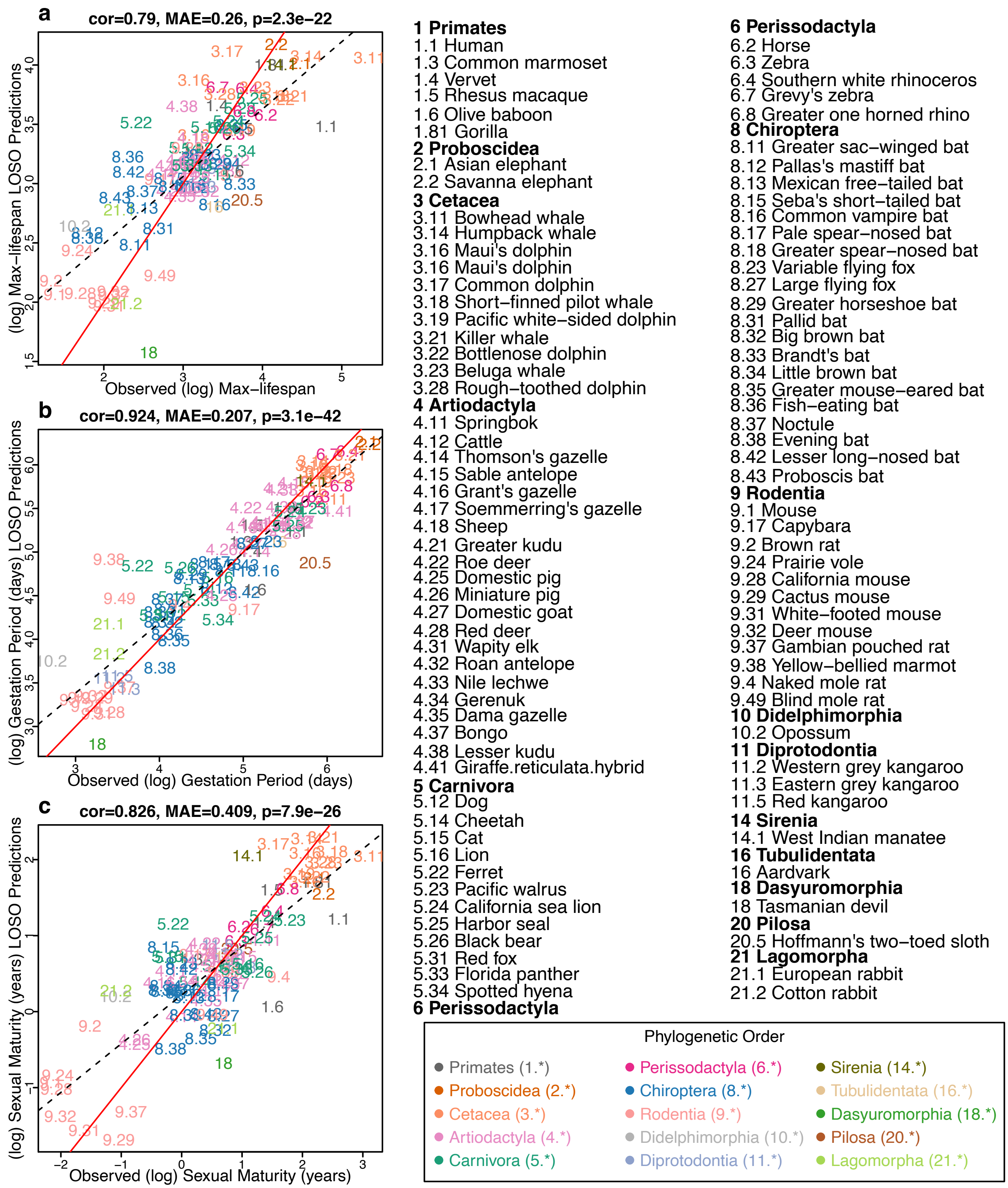


**Extended Data Fig. S2 | ElasticNet Predictor Based on Young Samples.**
Elastic Net Predictor, Leave One Species Out analysis, fitted on a subset of all young samples (n=99). Young samples are defined as a sample's age is younger than species average age at sex maturation. Feature filtering and Elastic Net tuning parameter set-up is the same as those for **Extended Data Fig.**

**S1**. Three panels show predictors for **a,** maximum lifespan (in years), **b,** gestation time (in days), and **c,** age at sexual maturity (in years). As with the main Figure 1, species appear as designated numbers in scatter plot panels; the corresponding common names and phylogenetic orders are annotated in figure legends; as indicated by the phylogenetic order legend, the whole number (number before the decimal separator) part of each mammalian number is assigned in accordance to the corresponding phylogenetic order.

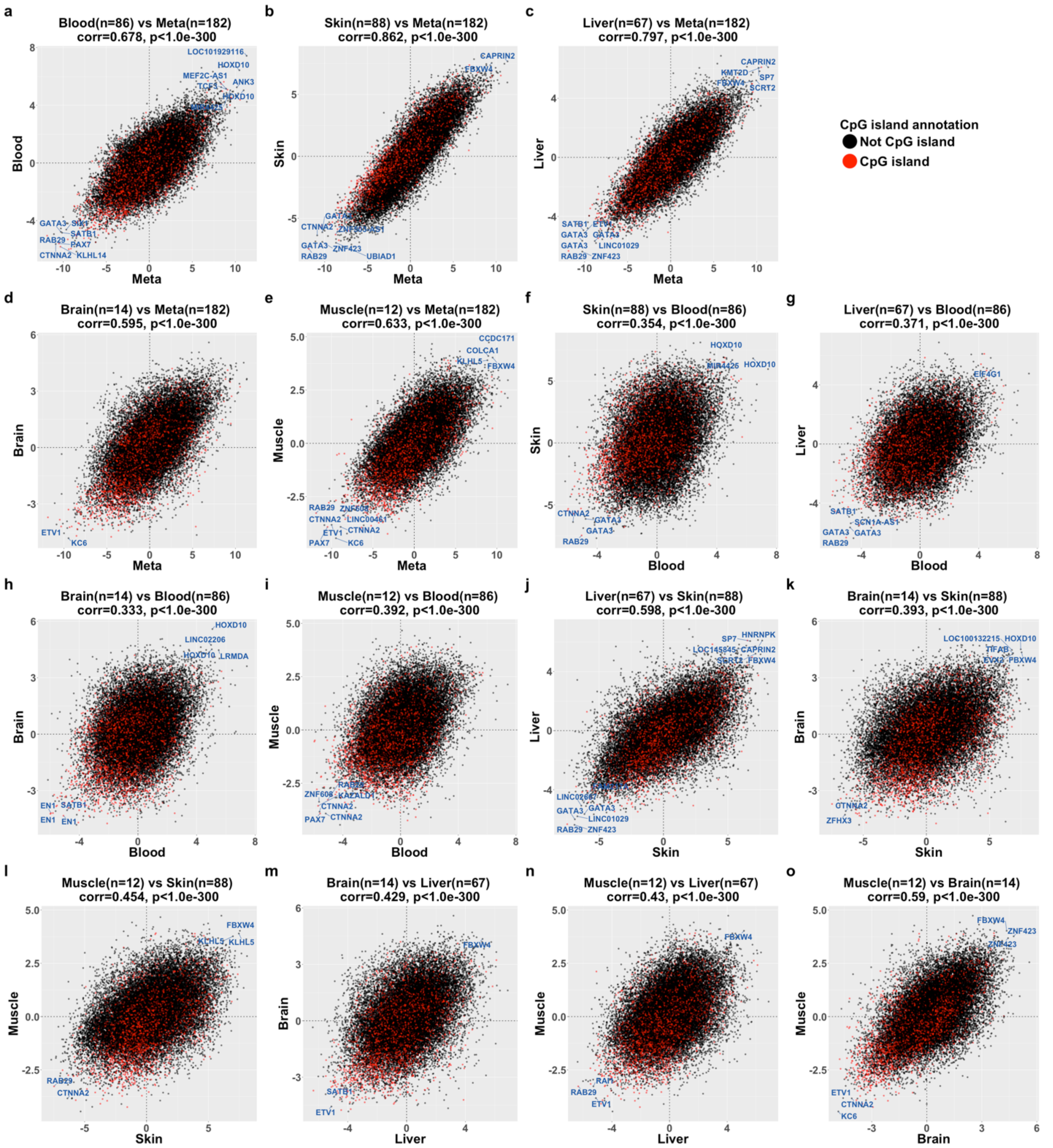


**Extended Data Fig. S3 | Generic Lifespan EWAS in different tissues from Eutherian species.** Scatter plot of CpG Z statistics agreements between tissues, stratified by CpG island annotations (not island: black, island: red). Both x-and-y-axes are CpG Z statistics for all CpG probes considered mappable to humans and mice (27k), but from different EWAS analyses. Panels show agreements between **a,** blood vs. meta, **b,** skin vs. meta, **c,** liver vs. meta, **d,** brain vs. meta, **e,** muscle vs. meta, **f,** skin vs. blood, **g,** liver vs. blood, **h,** brain vs. blood, i, muscle vs. blood, **j,** liver vs. skin, **k,** brain vs. skin, **l,** muscle vs. brain, **m,** brain vs. liver, **n,** muscle vs. liver, **o,** muscle vs. brain.

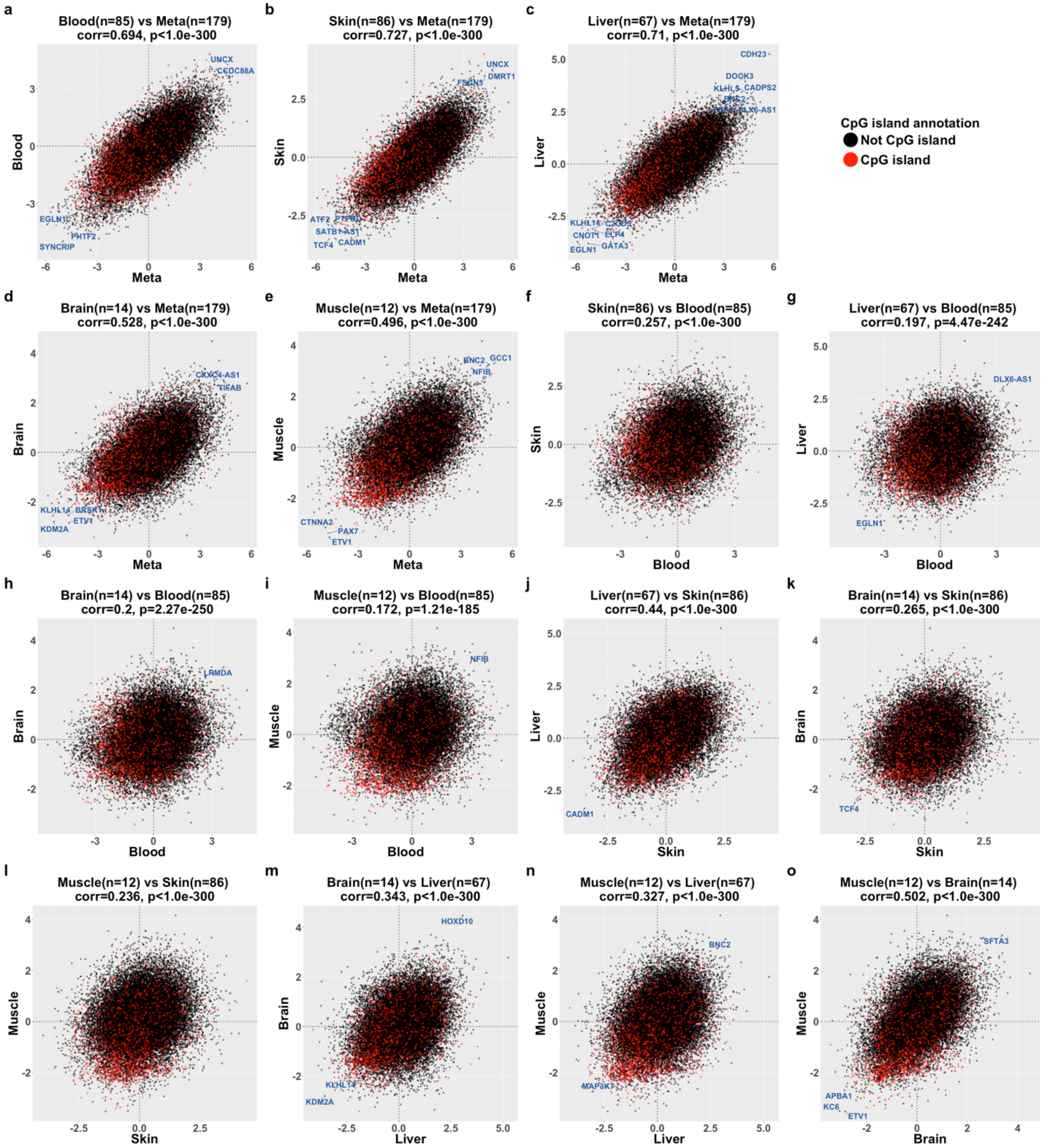


**Extended Data Fig. S4 | Phylogenetic EWAS agreement in various tissues, Eutherians.** Scatter plot of CpG Z statistics between tissues, stratified by CpG island annotations (not island: black, island:

red). Agreements are defined identically to **Extended Data Fig. S3.** Panels show agreements between **a,** blood vs. meta, **b,** skin vs. meta, **c,** liver vs. meta, **d,** brain vs. meta, **e,** muscle vs. meta, **f,** skin vs. blood, **g,** liver vs. blood, **h,** brain vs. blood, i, muscle vs. blood, **j,** liver vs. skin, **k,** brain vs. skin, **l,** muscle vs. brain, **m,** brain vs. liver, **n,** muscle vs. liver, **o,** muscle vs. brain.

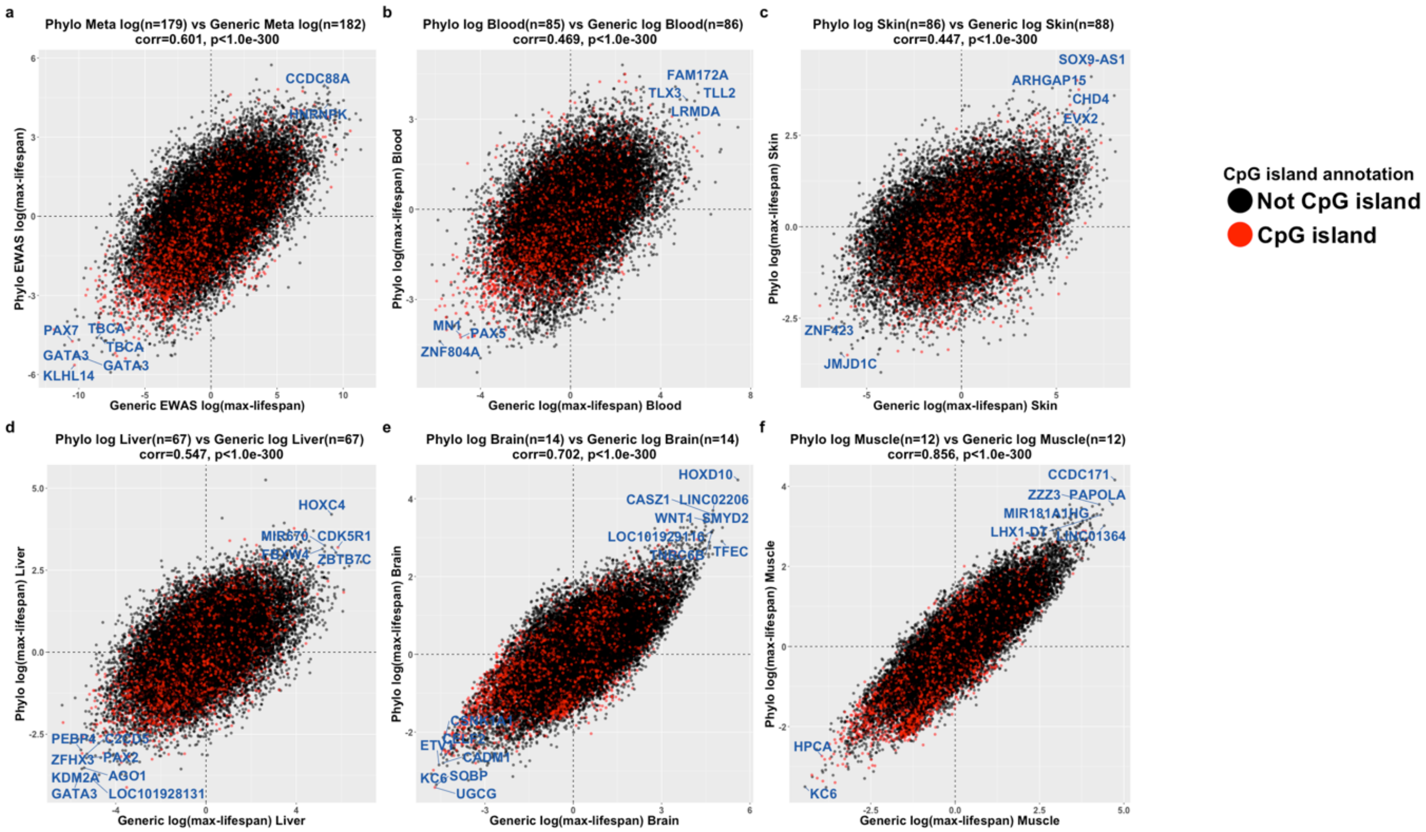


**Extended Data Fig. S5 | Simple Linear Regression (Generic) and phylogenetic Regression EWAS agreement.** Scatter plot of CpG Z statistics across phylogenetic Generic EWAS vs. Phylogenetic EWAS. Similar to **Extended Data Fig. S2**, panel titles and axes labels indicate agreements between EWAS analyses. Panels show agreements between, **a** meta phylogenetic vs. generic EWAS, **b,** phylogenetic vs. generic EWAS in blood, **c,** phylogenetic vs. generic EWAS in skin, **d,** phylogenetic vs. generic EWAS in liver, **e,** phylogenetic vs. generic EWAS in brain, **f,** phylogenetic vs. generic EWAS in muscle.

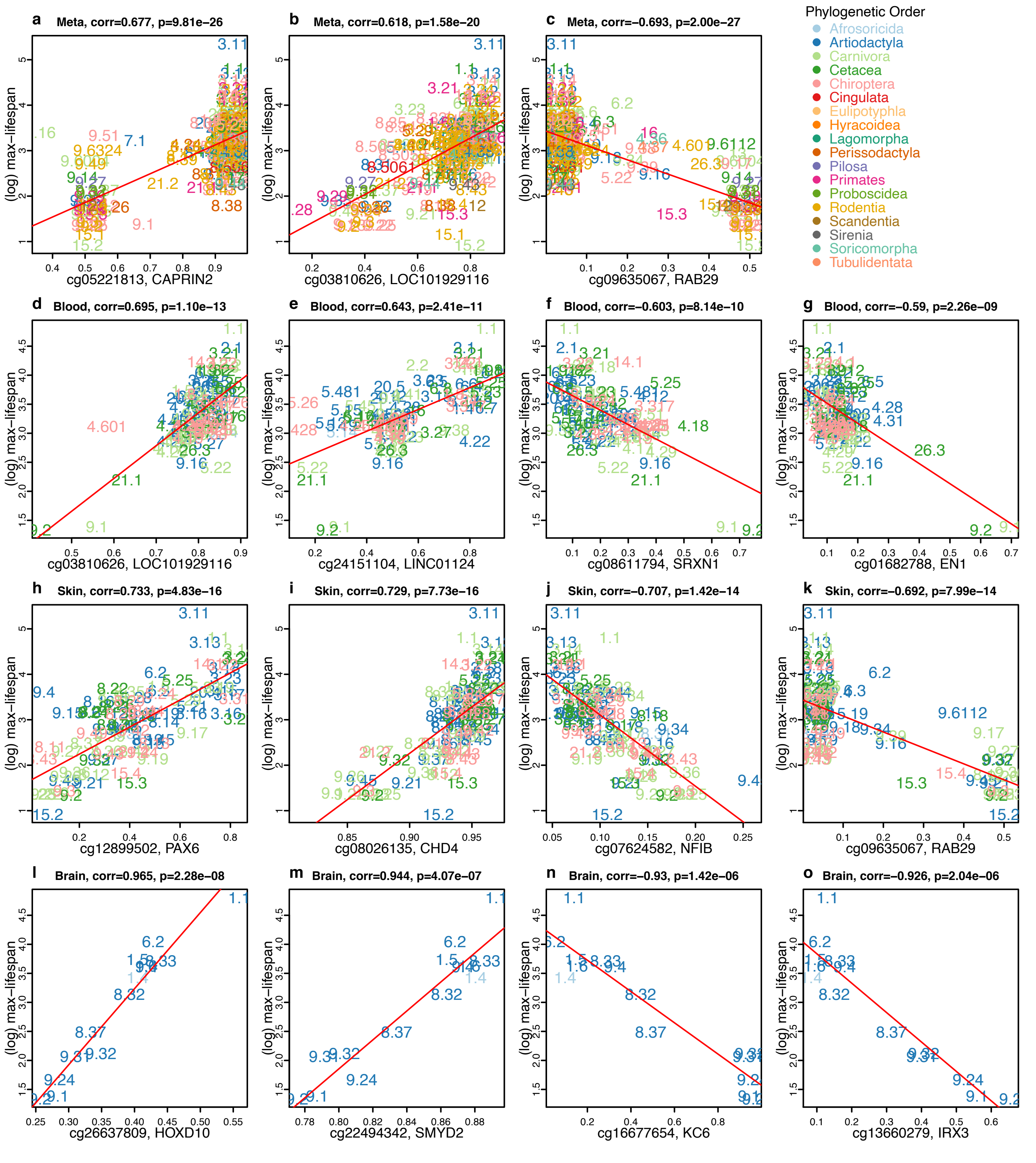


**Extended Data Fig. S6 | Top Significant CpG sites Scatter Plot, Eutherians.** Scatter plot of selected CpG probes from EWAS fitted on maximum lifespan. Each panel reports the Pearson correlation and student t.test pvalue. Stouffer method was used to summarize Z statistics from various tissues as a meta-analysis (Table 2.1.1). Panels show scatter plots of top 3 CpGs from **a-c,** meta-analysis for tissues, **b-g,** top 4 CpG in from blood tissues, **h-k,** top 4 CpGs from skin tissues, **l-o,** top 4 CpGs from brain tissues.

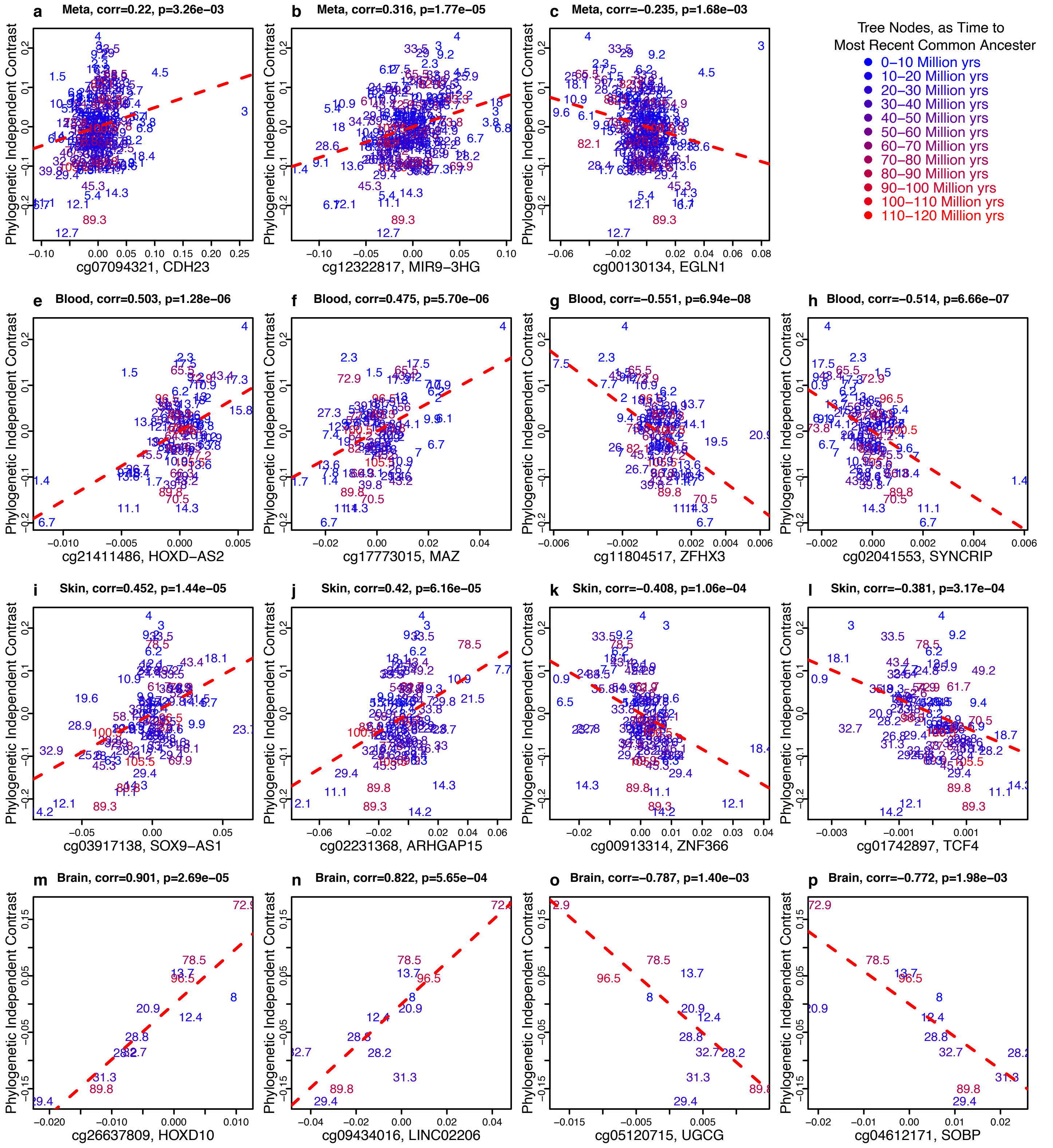


**Supplementary Figure S7 | Top Significant CpG sites phylogenetic independent contrast plot, Eutherians.** Scatter plot of CpG methylation and maximum lifespan, transformed and scaled to phylogenetic independent contrasts, based on all available samples. Each panel reports the Pearson correlation and student t.test pvalue. Stouffer method was used to summarize Z statistics from various tissues as a meta-analysis (Table 2.2.1). In order to properly visualize sample correlations, phylogenetic independent contrast plots select parent nodes that are of relatively similar distances to each other [1]. We color coded these common ancestor nodes as time to present, in millions of years. Panels show scatter plots of top 3 CpGs from **a-c,** meta analysis for tissues, **b-g,** top 4 CpG in from blood tissues, **h-k,** top 4 CpGs from skin tissues, **l-o,** top 4 CpGs from brain tissues

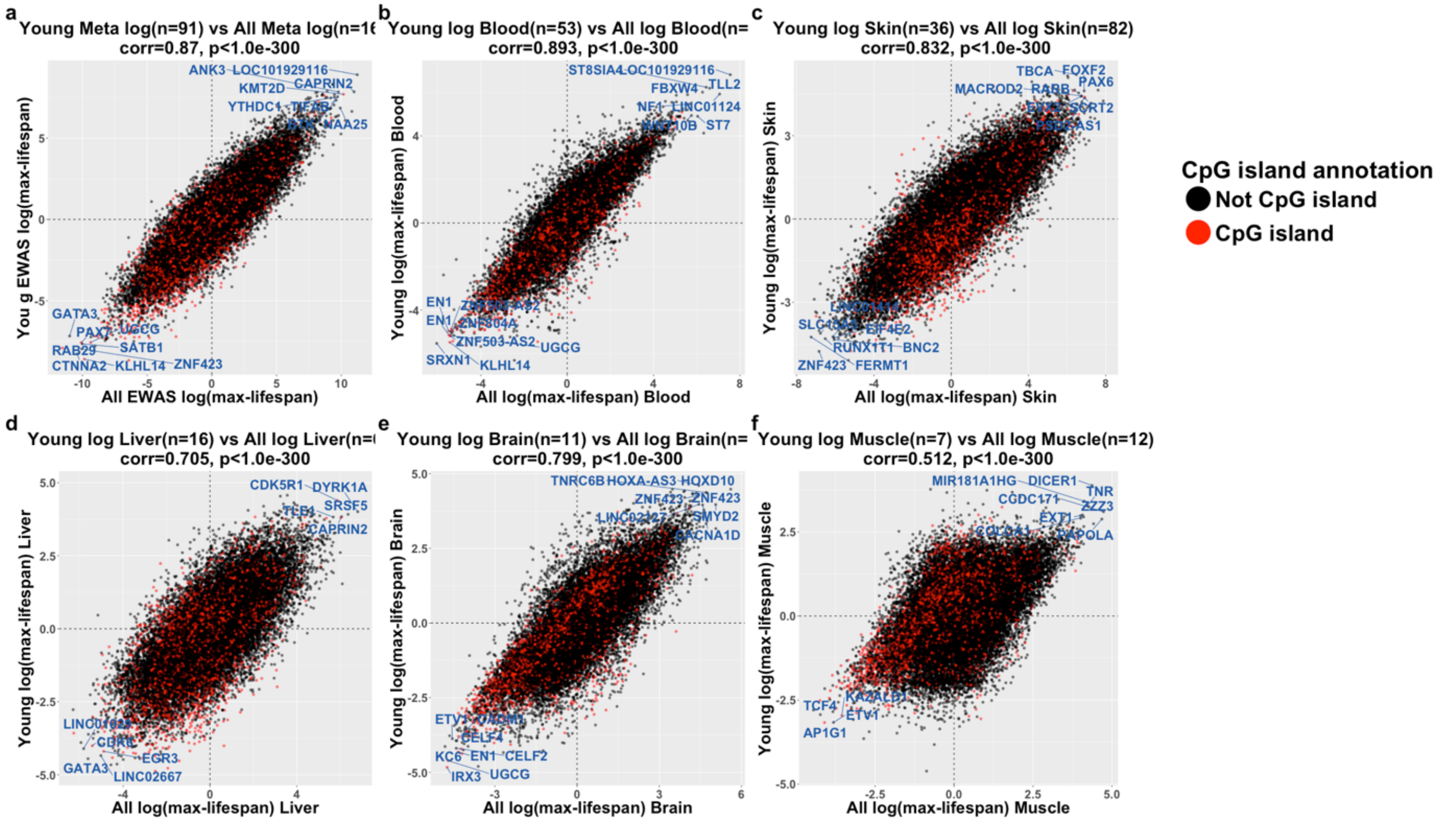


**Supplementary Figure S8 | Generic EWAS agreements between all samples and samples younger than 3 years of age.** Similar to Sup. Fig. S4 and S5, but agreements between EWAS based on samples younger than 3 years of age only EWAS based on all available samples. Panels show agreements between, **a** meta phylogenetic vs. generic EWAS, **b,** phylogenetic vs. generic EWAS in blood, **c,** phylogenetic vs. generic EWAS in skin, **d,** phylogenetic vs. generic EWAS in liver, **e,** phylogenetic vs. generic EWAS in brain, **f,** phylogenetic vs. generic EWAS in muscle.

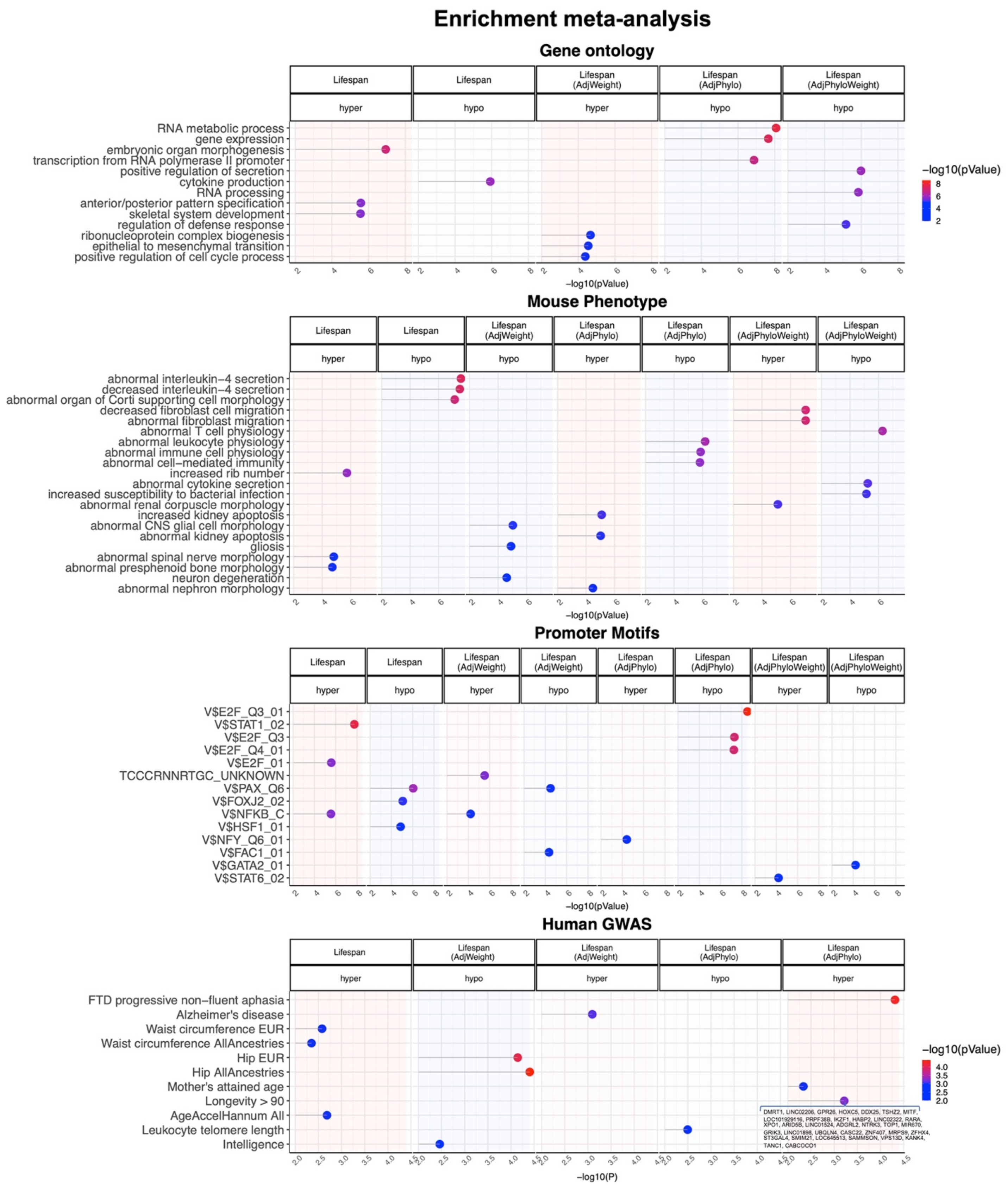
Enrichment meta-analysis
Gene ontology
Lifespan hyper
Lifespan hypo
Lifespan (AdjWeight) hyper
Lifespan (AdjPhylo) hypo
Lifespan (AdjPhyloWeight) hypo
RNA metabolic process
gene expression
embryonic organ morphogenesis
transcription from RNA polymerase II promoter
positive regulation of secretion
cytokine production
RNA processing
anterior/posterior pattern specification
skeletal system development
regulation of defense response
ribonucleoprotein complex biogenesis
epithelial to mesenchymal transition
positive regulation of cell cycle process
−log10(pValue)
Mouse Phenotype
Lifespan hyper
Lifespan hypo
Lifespan (AdjWeight) hypo
Lifespan (AdjPhylo) hyper
Lifespan (AdjPhylo) hypo
Lifespan (AdjPhyloWeight) hyper
Lifespan (AdjPhyloWeight) hypo
abnormal interleukin−4 secretion
decreased interleukin−4 secretion
abnormal organ of Corti supporting cell morphology
decreased fibroblast cell migration
abnormal fibroblast migration
abnormal T cell physiology
abnormal leukocyte physiology
abnormal immune cell physiology
abnormal cell−mediated immunity
increased rib number
abnormal cytokine secretion
increased susceptibility to bacterial infection
abnormal renal corpuscle morphology
increased kidney apoptosis
abnormal CNS glial cell morphology
abnormal kidney apoptosis
gliosis
abnormal spinal nerve morphology
abnormal presphenoid bone morphology
neuron degeneration
abnormal nephron morphology
−log10(pValue)
Promoter Motifs
Lifespan hyper
Lifespan hypo
Lifespan (AdjWeight) hyper
Lifespan (AdjWeight) hypo
Lifespan (AdjPhylo) hyper
Lifespan (AdjPhylo) hypo
Lifespan (AdjPhyloWeight) hyper
Lifespan (AdjPhyloWeight) hypo
V$E2F_Q3_01
V$STAT1_02
V$E2F_Q3
V$E2F_Q4_01
V$E2F_01
TCCCRNNRTGC_UNKNOWN
V$PAX_Q6
V$FOXJ2_02
V$NFKB_C
V$HSF1_01
V$NFY_Q6_01
V$FAC1_01
V$GATA2_01
V$STAT6_02
−log10(pValue)
Human GWAS
Lifespan hyper
Lifespan (AdjWeight) hypo
Lifespan (AdjWeight) hyper
Lifespan (AdjPhylo) hypo
Lifespan (AdjPhylo) hyper
FTD progressive non−fluent aphasia
Alzheimer's disease
Waist circumference EUR
Waist circumference AllAncestries
Hip EUR
Hip AllAncestries
Mother's attained age
Longevity > 90
AgeAccelHannum All
Leukocyte telomere length
Intelligence
−log10(P)
−log10(pValue)

**Extended Data Fig. S9 | Gene set enrichment analysis of significant CpGs related to mammalian maximum lifespan.** The gene level enrichment was done using GREAT analysis using human background. The background probes were limited to 27,966 probes that were experimentally validated to work in both mouse and human genomes. Human GWAS enrichment was calculated by a hypergeometric test of the top 5% genomic regions involved in GWAS of complex traits-associated genes with the top lifespan related gene regions in our analysis. The biological processes were reduced to parent ontology terms using the "rrvgo" package. Input, top 500 CpGs per direction for each model. In each panel, the columns with no significant terms were removed to simplify the figure.

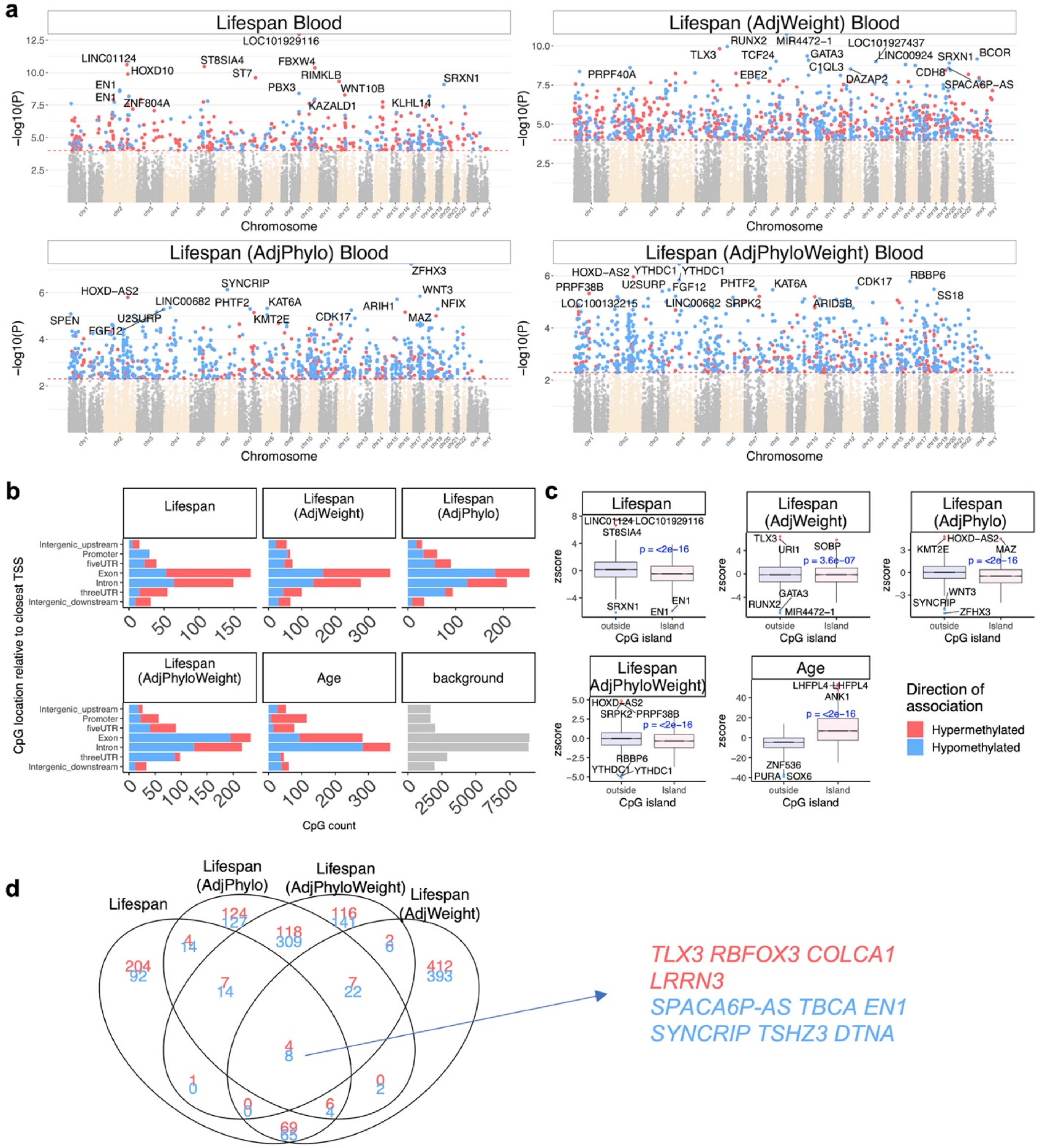


**Extended Data Fig. S10** | ***EWAS of mammalian maximum lifespan in blood.*** The associations were examined with four different models: 1) lifespan: each species as a datapoint in the model regardless of evolutionary distance. 2) lifespan adjusted for average species weight. 3) lifespan adjusted for evolutionary distance by phylogenetic regression. The evolutionary tree was acquired from TimeTree database [2]. 4) lifespan adjusted for both average adult species weight and evolutionary distance. Panel **a,** Manhattan plots of EWAS of maximum lifespan in 27966 probes that were experimentally validated to work in both mouse and human genomes. The coordinates are based on the alignment to the Human hg38 genome. The direction of associations with $p < 10^{-4}$ (red dotted line) is highlighted by red (hypermethylated) and blue (hypomethylated) colors. The phylogenetic regression models were studied at $p<0.005$. The top 15 CpGs were labeled by the neighboring genes, **b,** Location of top CpGs in each tissue relative to the closest transcriptional start site. A panel for the top 1000 age related CpGs was added to the figure for comparison. The grey color in the last panel represents the location of 27966 mammalian BeadChip array probes mapped to the human hg38 genome, **c,** Boxplot of association with mammalian maximum lifespan by human CpG island status. The mean difference was tested by student T test. A panel for the top 1000 age related CpGs was added to the figure for comparison, **d,** Venn diagram of the overlap in the top 1000 (500 per direction) significant CpGs for different models of EWAS of lifespan. The overlap hits were labeled with its neighboring gene in the plot.

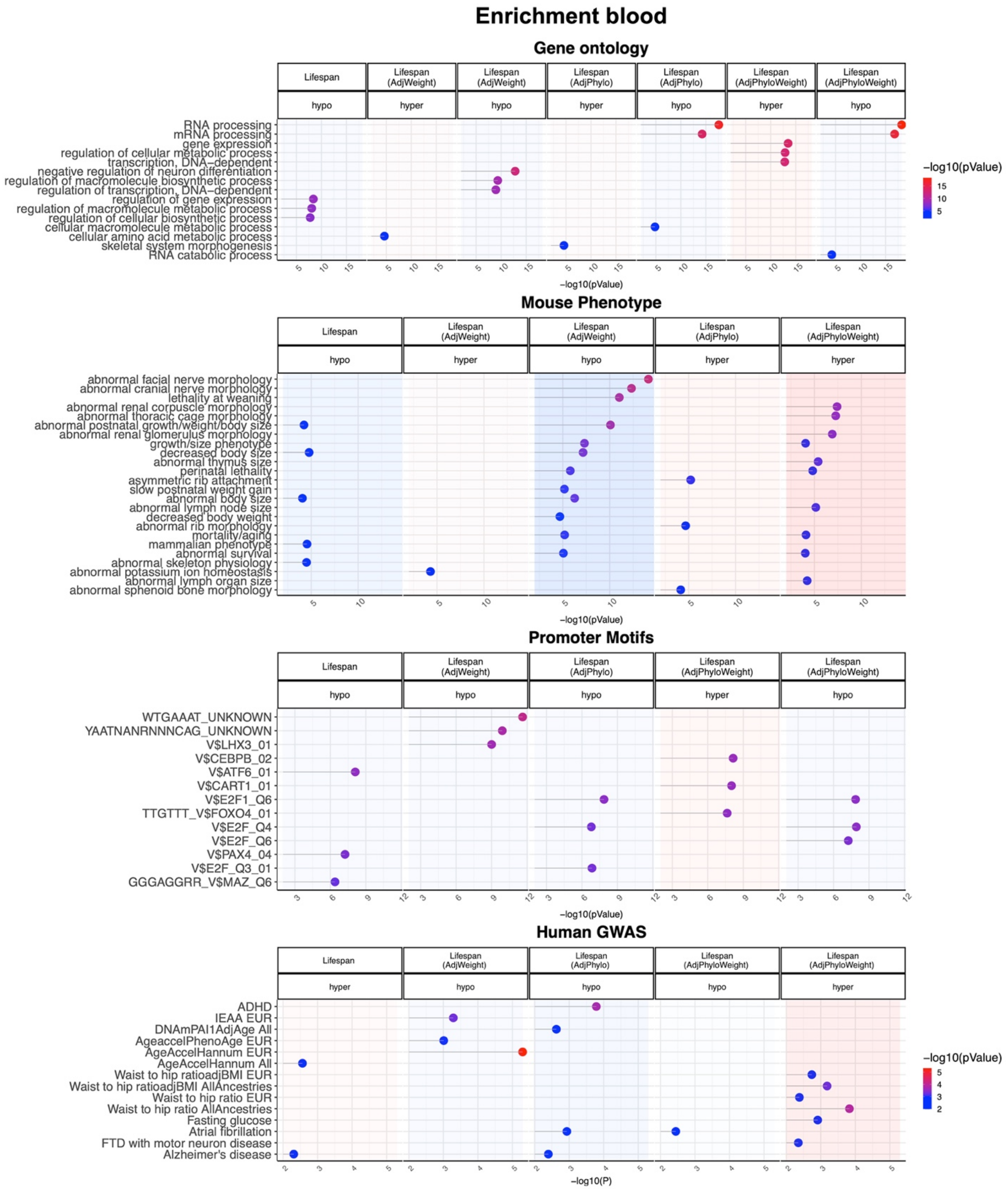
Enrichment blood
Gene ontology
Lifespan
Lifespan (AdjWeight)
Lifespan (AdjPhylo)
Lifespan (AdjPhyloWeight)
hypo
hyper
RNA processing
mRNA processing
gene expression
regulation of cellular metabolic process
transcription, DNA−dependent
negative regulation of neuron differentiation
regulation of macromolecule biosynthetic process
regulation of transcription, DNA−dependent
regulation of gene expression
regulation of macromolecule metabolic process
regulation of cellular biosynthetic process
cellular macromolecule metabolic process
cellular amino acid metabolic process
skeletal system morphogenesis
RNA catabolic process
−log10(pValue)
Mouse Phenotype
abnormal facial nerve morphology
abnormal cranial nerve morphology
lethality at weaning
abnormal renal corpuscle morphology
abnormal thoracic cage morphology
abnormal postnatal growth/weight/body size
abnormal renal glomerulus morphology
growth/size phenotype
decreased body size
abnormal thymus size
perinatal lethality
asymmetric rib attachment
slow postnatal weight gain
abnormal body size
abnormal lymph node size
decreased body weight
abnormal rib morphology
mortality/aging
mammalian phenotype
abnormal survival
abnormal skeleton physiology
abnormal potassium ion homeostasis
abnormal lymph organ size
abnormal sphenoid bone morphology
Promoter Motifs
WTGAAAT_UNKNOWN
YAATNANRNNNCAG_UNKNOWN
V$LHX3_01
V$CEBPB_02
V$ATF6_01
V$CART1_01
V$E2F1_Q6
TTGTTT_V$FOXO4_01
V$E2F_Q4
V$E2F_Q6
V$PAX4_04
V$E2F_Q3_01
GGGAGGRR_V$MAZ_Q6
Human GWAS
ADHD
IEAA EUR
DNAmPAI1AdjAge All
AgeaccelPhenoAge EUR
AgeAccelHannum EUR
AgeAccelHannum All
Waist to hip ratioadjBMI EUR
Waist to hip ratioadjBMI AllAncestries
Waist to hip ratio EUR
Waist to hip ratio AllAncestries
Fasting glucose
Atrial fibrillation
FTD with motor neuron disease
Alzheimer's disease
−log10(P)

**Extended Data Fig. S11 | Gene set enrichment analysis of significant CpGs related to mammalian maximum lifespan in blood.** The gene level enrichment was done using GREAT analysis using human background. The background probes were limited to 27,966 probes that were experimentally validated to work in both mouse and human genomes. Human GWAS enrichment was calculated by a hypergeometric test of the top 5% genomic regions involved in GWAS of complex traits-associated genes with the top lifespan related gene regions in our analysis. The biological processes were reduced to parent ontology terms using the “rrvgo” package. Input: Lifespan hypo/hyper, 197/295 CpGs; Lifespan (AdjWeight) hypo/hyper, 500/500; Lifespan (AdjPhylo) hypo/hyper, 500/270; Lifespan (AdjPhyloWeight) hypo/hyper, 500/255. In each panel, the columns with no significant terms were removed to simplify the figure.

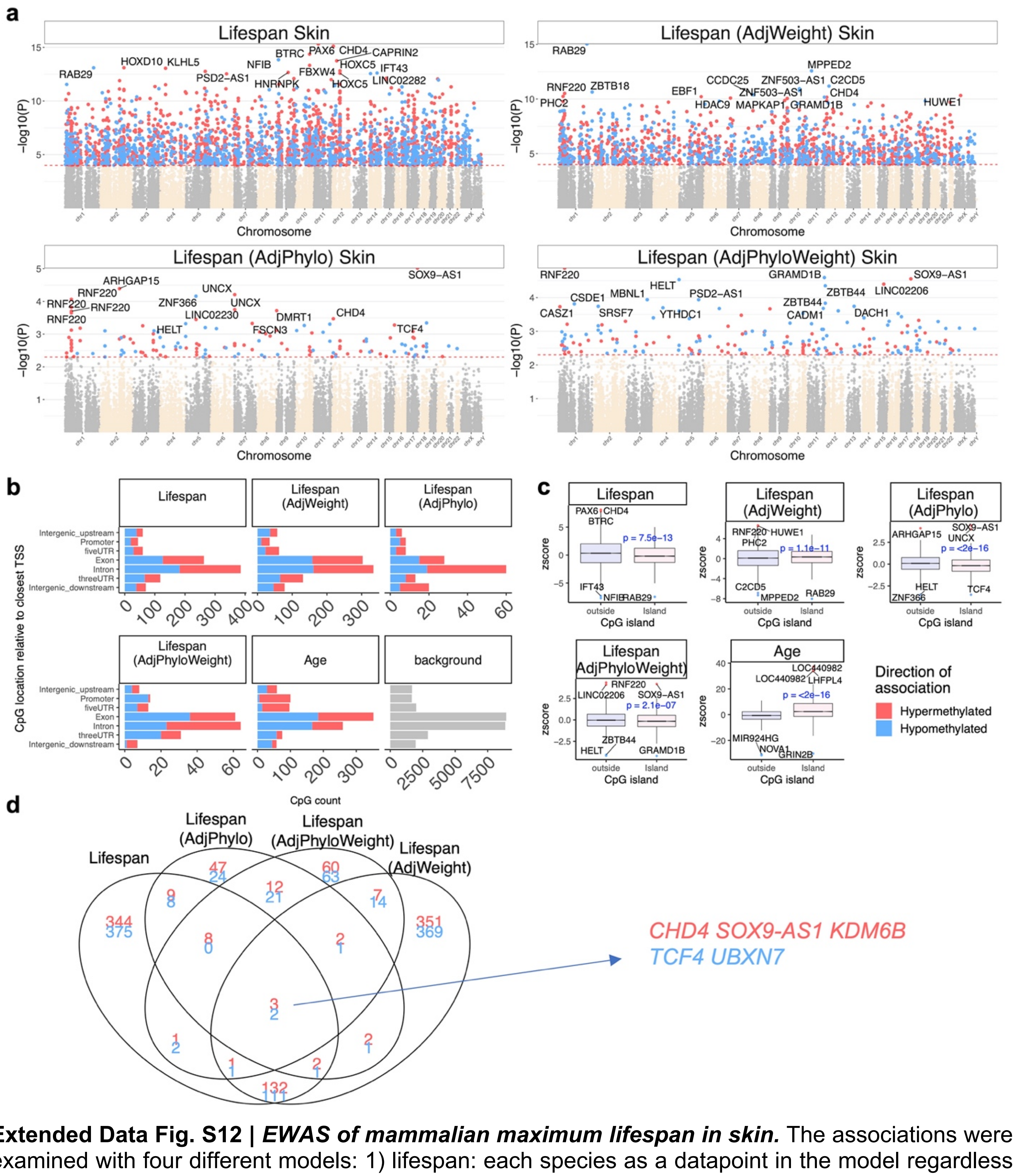

**Extended Data Fig. S12 | *EWAS of mammalian maximum lifespan in skin.*** The associations were examined with four different models: 1) lifespan: each species as a datapoint in the model regardless of evolutionary distance. 2) lifespan adjusted for average species weight. 3) lifespan adjusted for evolutionary distance by phylogenetic regression. The evolutionary tree was acquired from TimeTree database [2]. 4) lifespan adjusted for both average adult species weight and evolutionary distance. Panel **a,** Manhattan plots of EWAS of maximum lifespan in 27966 probes that were experimentally validated to work in both mouse and human genomes. The coordinates are based on the alignment to the Human hg38 genome. The direction of associations with $p < 10^{-4}$ (red dotted line) is highlighted by red (hypermethylated) and blue (hypomethylated) colors. The phylogenetic regression models were studied at $p < 0.005$. The top 15 CpGs were labeled by the neighboring genes, **b,** Location of top CpGs in each tissue relative to the closest transcriptional start site. A panel for the top 1000 age related CpGs was added to the figure for comparison. The grey color in the last panel represents the location of 27966 mammalian BeadChip array probes mapped to the human hg38 genome, **c,** Boxplot of association with mammalian maximum lifespan by human CpG island status. The mean difference was tested by student T test. A panel for the top 1000 age related CpGs was added to the figure for comparison, **d** Venn diagram of the overlap in the top 1000 (500 per direction) significant CpGs for different models of EWAS of lifespan. The overlap hits were labeled with its neighboring gene in the plot.

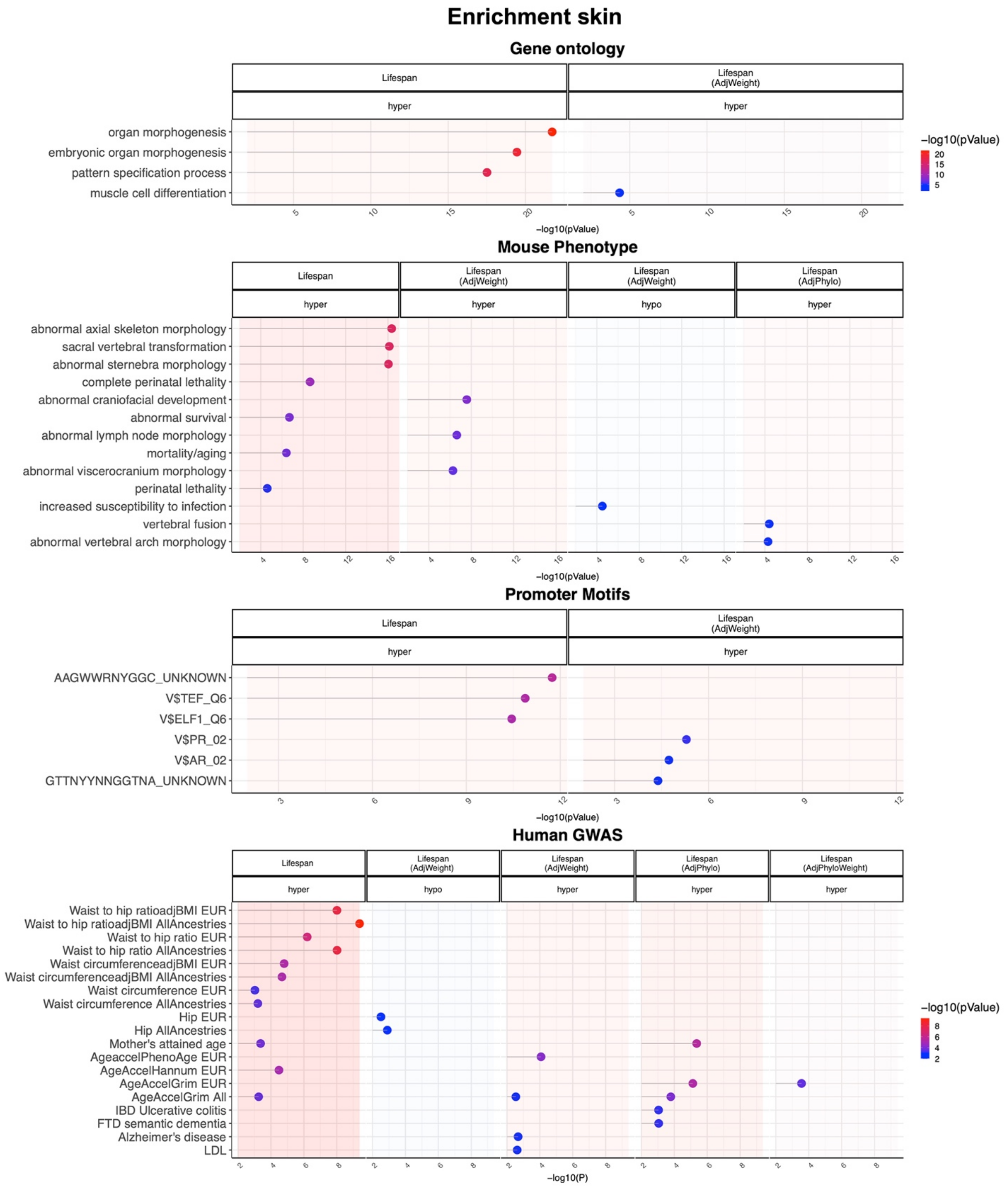

Enrichment skin
Gene ontology
Lifespan
hyper
Lifespan (AdjWeight)
hyper
organ morphogenesis
embryonic organ morphogenesis
pattern specification process
muscle cell differentiation
5
10
15
20
−log10(pValue)
−log10(pValue)
20
15
10
5
Mouse Phenotype
Lifespan
hyper
Lifespan (AdjWeight)
hyper
Lifespan (AdjWeight)
hypo
Lifespan (AdjPhylo)
hyper
abnormal axial skeleton morphology
sacral vertebral transformation
abnormal sternebra morphology
complete perinatal lethality
abnormal craniofacial development
abnormal survival
abnormal lymph node morphology
mortality/aging
abnormal viscerocranium morphology
perinatal lethality
increased susceptibility to infection
vertebral fusion
abnormal vertebral arch morphology
4
8
12
16
−log10(pValue)
Promoter Motifs
Lifespan
hyper
Lifespan (AdjWeight)
hyper
AAGWWRNYGGC_UNKNOWN
V$TEF_Q6
V$ELF1_Q6
V$PR_02
V$AR_02
GTTNYYNNGGTNA_UNKNOWN
3
6
9
12
−log10(pValue)
Human GWAS
Lifespan
hyper
Lifespan (AdjWeight)
hypo
Lifespan (AdjWeight)
hyper
Lifespan (AdjPhylo)
hyper
Lifespan (AdjPhyloWeight)
hyper
Waist to hip ratioadjBMI EUR
Waist to hip ratioadjBMI AllAncestries
Waist to hip ratio EUR
Waist to hip ratio AllAncestries
Waist circumferenceadjBMI EUR
Waist circumferenceadjBMI AllAncestries
Waist circumference EUR
Waist circumference AllAncestries
Hip EUR
Hip AllAncestries
Mother's attained age
AgeaccelPhenoAge EUR
AgeAccelHannum EUR
AgeAccelGrim EUR
AgeAccelGrim All
IBD Ulcerative colitis
FTD semantic dementia
Alzheimer's disease
LDL
2
4
6
8
−log10(P)
−log10(pValue)
8
6
4
2

**Extended Data Fig. S13 | Gene set enrichment analysis of significant CpGs related to mammalian maximum lifespan in skin.** The gene level enrichment was done using GREAT analysis using human background. The background probes were limited to 27,966 probes that were experimentally validated to work in both mouse and human genomes. Human GWAS enrichment was calculated by a hypergeometric test of the top 5% genomic regions involved in GWAS of complex traits-associated genes with the top lifespan related gene regions in our analysis. The biological processes were reduced to parent ontology terms using the “rrvgo” package. Input: Lifespan hypo/hyper, 500/500 CpGs; Lifespan (AdjWeight) hypo/hyper, 500/500; Lifespan (AdjPhylo) hypo/hyper, 58/85; Lifespan (AdjPhyloWeight) hypo/hyper, 104/94. In each panel, the columns with no significant terms were removed to simplify the figure.

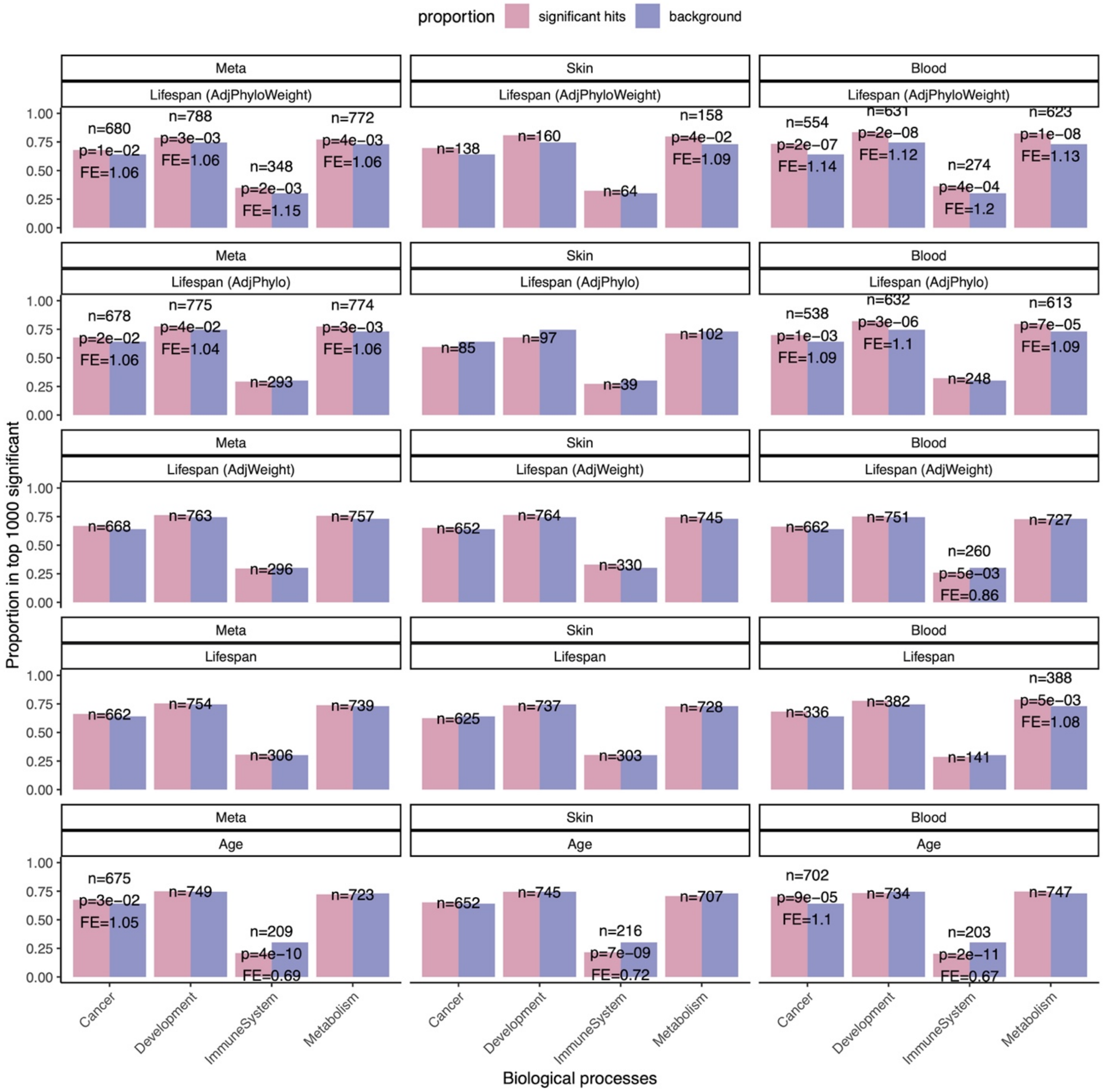


**Extended Data Fig. S14 | Biological processes of genes adjacent to the CpGs associated with mammalian maximum lifespan.** The biological processes are defined by text mining of selected keywords from the Gene-ontology terms related to each gene. Keywords: Metabolism (metabolism, synthesis, catabolism, anabolism, biosynthesis), Development (growth, development, morphology, differentiate, differentiation, stem cell), Immune system (cytokine, IL, chemokine, immune, antigen, antibody, inflammation, stress, shock, microbe, virus), Cancer (cancer, tumor, neoplasm, sarcoma, carcinoma, leukemia). The columns with significant proportion difference from the background are labeled with p-value, and fold enrichment (FE). The number of significant probes for each biological process are labeled in the figure. Blood input: Lifespan hypo/hyper, 197/295 CpGs; Lifespan (AdjWeight) hypo/hyper, 500/500; Lifespan (AdjPhylo) hypo/hyper, 500/270; Lifespan (AdjPhyloWeight) hypo/hyper, 500/255. Skin input: Lifespan hypo/hyper, 500/500 CpGs; Lifespan (AdjWeight) hypo/hyper, 500/500; Lifespan (AdjPhylo) hypo/hyper, 58/85; Lifespan (AdjPhyloWeight) hypo/hyper, 104/94. Lifespan Meta input: Lifespan hypo/hyper, 500/500 CpGs; Lifespan (AdjWeight) hypo/hyper, 500/500; Lifespan (AdjPhylo) hypo/hyper, 500/500; Lifespan (AdjPhyloWeight) hypo/hyper, 500/500; Age hypo/hyper, 500/500.

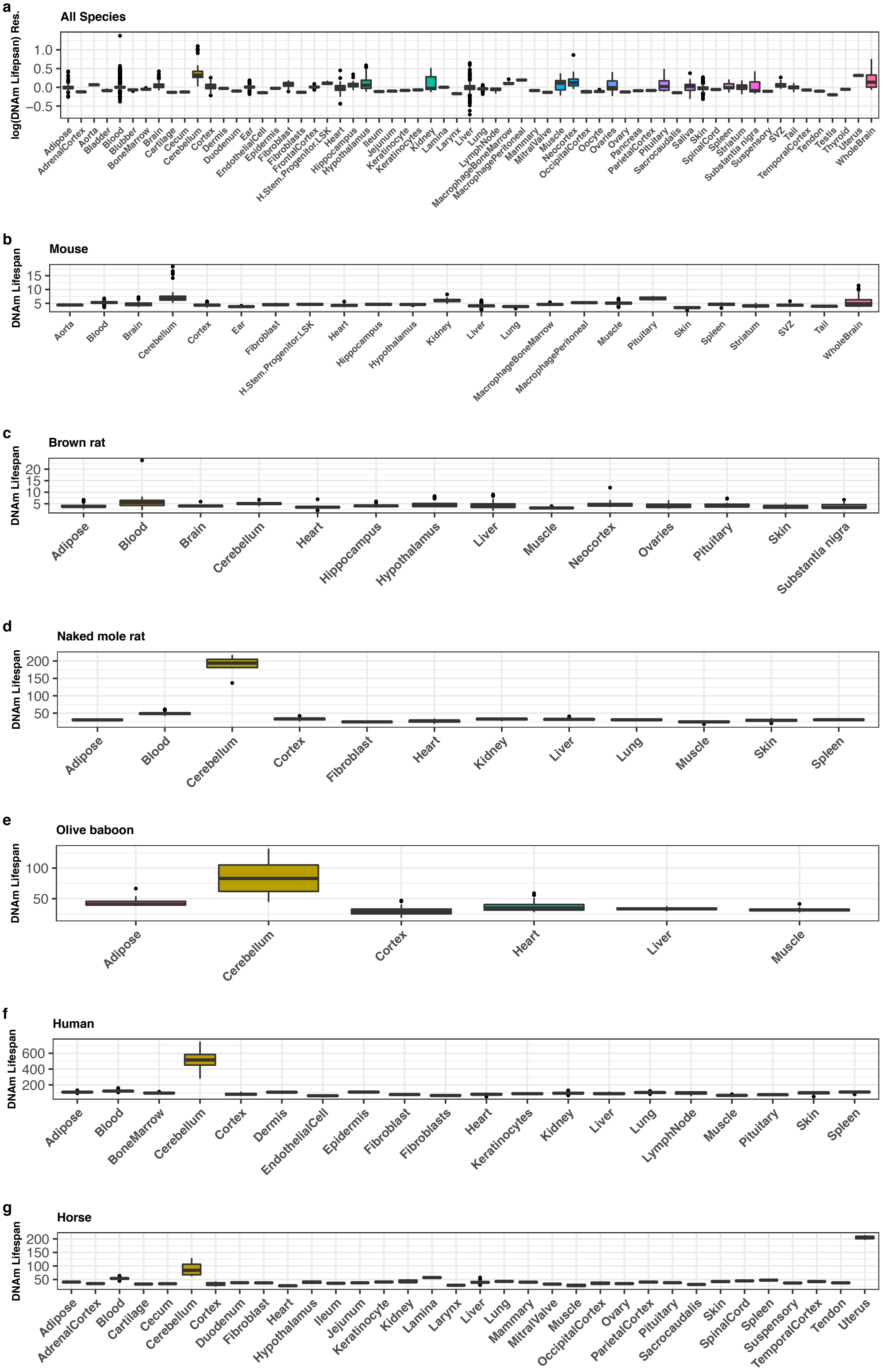
a
All Species
log(DNAm Lifespan) Res.
1.0
0.5
0.0
-0.5
Adipose AdrenalCortex Aorta Bladder Blood Blubber BoneMarrow Brain Cartilage Cecum Cerebellum Cortex Dermis Duodenum Ear EndothelialCell Epidermis Fibroblast Fibroblasts FrontalCortex H.Stem.Progenitor.LSK Heart Hippocampus Hypothalamus Ileum Jejunum Keratinocyte Keratinocytes Kidney Lamina Larynx Liver Lung LymphNode MacrophageBoneMarrow MacrophagePeritoneal Mammary MitralValve Muscle Neocortex OccipitalCortex Oocyte Ovaries Ovary Pancreas ParietalCortex Pituitary Sacrocaudalis Saliva Skin SpinalCord Spleen Striatum Substantia nigra Suspensory SVZ Tail TemporalCortex Tendon Testis Thyroid Uterus WholeBrain
b
Mouse
DNAm Lifespan
15
10
5
Aorta Blood Brain Cerebellum Cortex Ear Fibroblast H.Stem.Progenitor.LSK Heart Hippocampus Hypothalamus Kidney Liver Lung MacrophageBoneMarrow MacrophagePeritoneal Muscle Pituitary Skin Spleen Striatum SVZ Tail WholeBrain
c
Brown rat
DNAm Lifespan
20
15
10
5
Adipose Blood Brain Cerebellum Heart Hippocampus Hypothalamus Liver Muscle Neocortex Ovaries Pituitary Skin Substantia nigra
d
Naked mole rat
DNAm Lifespan
200
150
100
50
Adipose Blood Cerebellum Cortex Fibroblast Heart Kidney Liver Lung Muscle Skin Spleen
e
Olive baboon
DNAm Lifespan
100
50
Adipose Cerebellum Cortex Heart Liver Muscle
f
Human
DNAm Lifespan
600
400
200
Adipose Blood BoneMarrow Cerebellum Cortex Dermis EndothelialCell Epidermis Fibroblast Fibroblasts Heart Keratinocytes Kidney Liver Lung LymphNode Muscle Pituitary Skin Spleen
g
Horse
DNAm Lifespan
200
150
100
50
Adipose AdrenalCortex Blood Cartilage Cecum Cerebellum Cortex Duodenum Fibroblast Heart Hypothalamus Ileum Jejunum Keratinocyte Kidney Lamina Larynx Liver Lung Mammary MitralValve Muscle OccipitalCortex Ovary ParietalCortex Pituitary Sacrocaudalis Skin SpinalCord Spleen Suspensory TemporalCortex Tendon Uterus

**Extended Data Fig. S15 | Tissue groups differences in predicted mammalian maximum lifespan.** Mammalian maximum lifespan predictor, based on averaged species methylation, was used to predict individual sample lifespans. The predicted values are grouped by sample tissue annotations. Panel **a** shows predicted maximum lifespans (DNAm lifespan) standardized residuals (Res.) by tissue groups in all species and samples; in order to show viewable scales in different species, due to their drastically different lifespans, we evaluated residuals standardized by species (log of predicted maximum lifespan minus log of observed maximum lifespan, results from which are divided by log of observed maximum lifespan of the species to which the samples belong); panel **b – g** shows boxplots of predicted lifespans in original scales (DNAm lifespan) by tissue groups; due to the fact that within species comparisons require no re-scaling, predicted lifespans (in years) are shown in these panels; Tissue type "H.Stem.Progenitor.LSK" stands for "LSK Progenitor  Hematopoietic Stem cells"

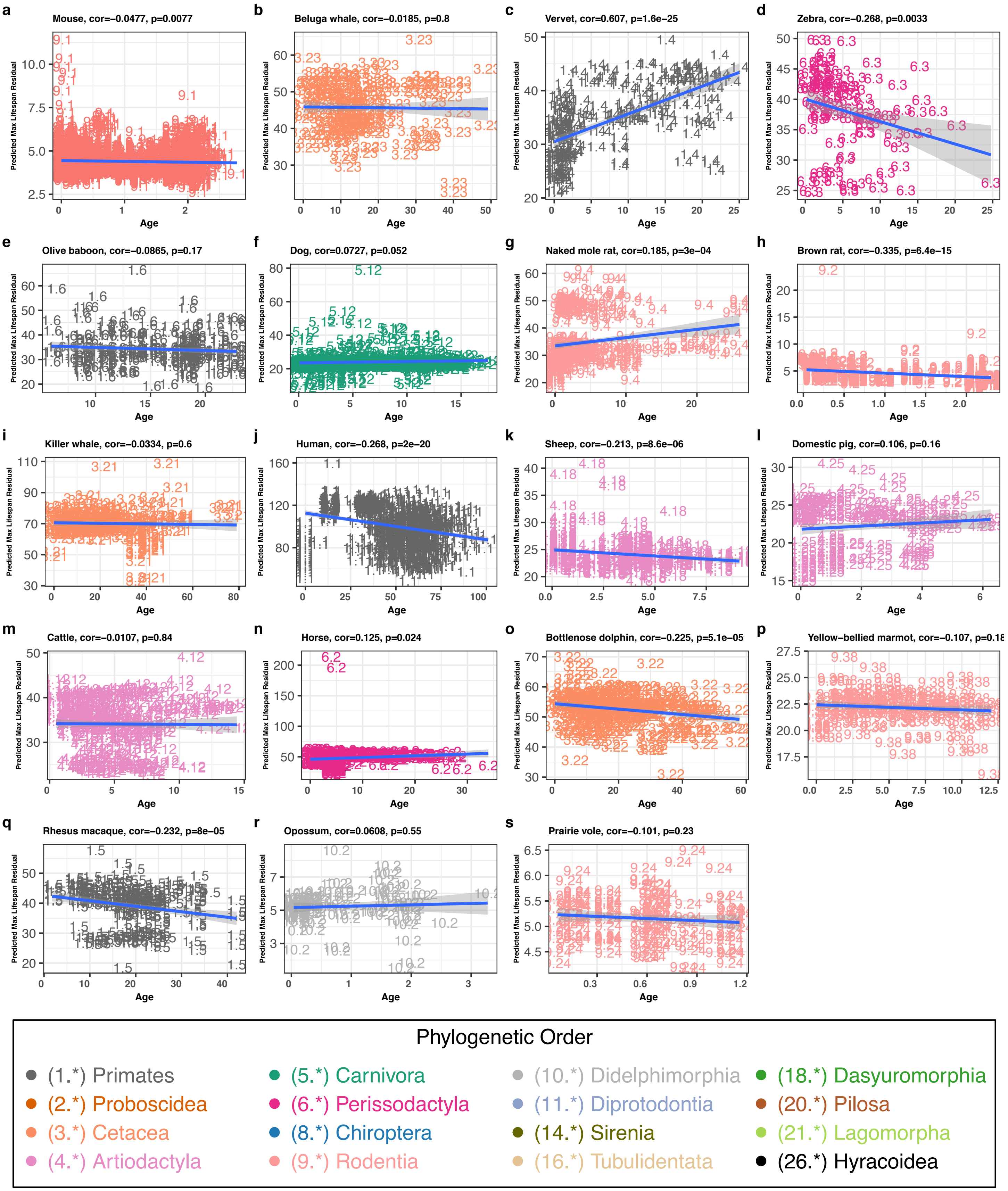
a
Mouse, cor=−0.0477, p=0.0077
b
Beluga whale, cor=−0.0185, p=0.8
c
Vervet, cor=0.607, p=1.6e−25
d
Zebra, cor=−0.268, p=0.0033
e
Olive baboon, cor=−0.0865, p=0.17
f
Dog, cor=0.0727, p=0.052
g
Naked mole rat, cor=0.185, p=3e−04
h
Brown rat, cor=−0.335, p=6.4e−15
i
Killer whale, cor=−0.0334, p=0.6
j
Human, cor=−0.268, p=2e−20
k
Sheep, cor=−0.213, p=8.6e−06
l
Domestic pig, cor=0.106, p=0.16
m
Cattle, cor=−0.0107, p=0.84
n
Horse, cor=0.125, p=0.024
o
Bottlenose dolphin, cor=−0.225, p=5.1e−05
p
Yellow−bellied marmot, cor=−0.107, p=0.18
q
Rhesus macaque, cor=−0.232, p=8e−05
r
Opossum, cor=0.0608, p=0.55
s
Prairie vole, cor=−0.101, p=0.23
Predicted Max Lifespan Residual
Age
Phylogenetic Order
(1.*) Primates
(2.*) Proboscidea
(3.*) Cetacea
(4.*) Artiodactyla
(5.*) Carnivora
(6.*) Perissodactyla
(8.*) Chiroptera
(9.*) Rodentia
(10.*) Didelphimorphia
(11.*) Diprotodontia
(14.*) Sirenia
(16.*) Tubulidentata
(18.*) Dasyuromorphia
(20.*) Pilosa
(21.*) Lagomorpha
(26.*) Hyracoidea

**Extended Data Fig. S16 | Correlation between maximum lifespan predictor and sample chronological ages.** Mammalian maximum lifespan predictor, based on averaged species methylation, was used to predict individual sample lifespans. The predicted values are also stratified by species. Only species with >100 sample sizes are shown. Each of **a-s** shows scatter plots of predicted lifespans in original scales vs. chronological age in specific species.
